# Unintended bias in the pursuit of collinearity solutions in fMRI analysis

**DOI:** 10.1101/2025.01.14.633053

**Authors:** Jeanette A. Mumford, Michael I. Demidenko, James M. Bjork, Bader Chaarani, Eric J. Feczko, Hugh P. Garavan, Donald J. Hagler, Steven M. Nelson, Tor D. Wager, Russell A. Poldrack

## Abstract

In task functional magnetic resonance imaging (fMRI), collinearity between task regressors in time series models may impact power. When collinearity is identified after data collection, researchers often modify the model in an effort to reduce collinearity. However, some model adjustments are suboptimal and may introduce bias into parameter estimates. Although relevant to many task-fMRI studies, we highlight these issues using the Monetary Incentive Delay (MID) task data from the Adolescent Brain Cognitive Development (ABCD^®^) study. We introduce a procedure to more directly quantify the impact of collinearity on task-relevant measures: a contrast-based variance inflation factor (cVIF). We also show that collinearity reduction strategies—such as omitting regressors for specific task components, using impulse regressors for extended activations, and ignoring response time variability—can bias contrast estimates. Finally, we present a “Saturated” model that includes all task components, including response times, aiming to reduce these biases while maintaining comparable levels of collinearity, as assessed by cVIF.

## 1 Introduction

Functional magnetic resonance imaging (fMRI) tasks often produce signals that are challenging to detect, especially when studying individual differences (M. I. Demidenko et al., 2024; Elliott et al., 2020; R. A. Poldrack et al., 2017). To address this, researchers prioritize models and study designs that maximize statistical power. Since collinearity between regressors can reduce power, efficiency optimization is often used prior to data collection to minimize collinearity (Friston et al., 1999; Liu et al., 2001). We assume the models used in design optimization and data analysis are identical to ensure the optimization results align as closely as possible to real data outcomes, aside from differences introduced by behavior-based or motion regressors that are unknown during design optimization. Certain tasks, like the Monetary Incentive Delay (MID) task, are inherently at risk for high collinearity. This arises because events that drive distinct neural processes are presented in rapid succession without intervening null events. As a result, the BOLD signals associated with these events overlap significantly, leaving minimal unique, non-overlapping signal to support the estimation of BOLD responses due to each event separately. Here, we will focus on the version of MID task used in the Adolescent Brain Cognitive Development (ABCD^®^) study, in which each trial consists of a cue, fixation, probe, and feedback presented consecutively, with no intervals in between. This type of design is likely to have substantial overlap between the BOLD signals emerging from adjacent task components within a trial. In such a design, a common practice to resolve collinearity is to omit regressors for a subset of trial components (e.g., fixation and probe in the MID task) or model the neural activity duration as an impulse. However, these common practices can bias the remaining estimates, making any analysis of “power benefit” due to collinearity reduction irrelevant. We examine these biases using the MID task data from the ABCD Study. Then, we illustrate the unintentional biases that occur as a result of collinearity avoidance strategies and propose a modeling strategy to reduce known biases, to the best of current knowledge, in the analysis of MID and other similarly designed fMRI tasks with closely spaced trial components.

This paper is divided into four parts. First, we review best practices for modeling fMRI data, focusing on how to assess collinearity, when it is acceptable and how to address it. Second, we introduce the MID task, its history, its design as implemented in the ABCD^®^ study and a brief overview of historical modeling decisions. Third, we demonstrate how modeling choices affect key contrast estimates for the MID task that are commonly used in the literature and are distributed by the ABCD consortium (Chaarani et al., 2021), using simulations to highlight potential biases from unmodeled neural activity and then apply these models to real fMRI data from the ABCD^®^ study’s MID task acquisition. Finally, we discuss our findings and provide recommendations. Although centered on the MID task, our modeling strategy is broadly applicable to task fMRI designs that involve multiple events in rapid succession.

### 1.1 fMRI time series modeling and collinearity

A core principle of fMRI modeling is to include all events expected to elicit neural activity. However, this becomes challenging when task designs cannot be modified, either due to inherent constraints or discoveries after data collection. Modeling each event individually can lead to high collinearity, forcing researchers to make compromises. Our goal is to balance this trade-off between contrast estimate variability, bias, and parameter interpretability. To support this, we introduce a new tool for estimating variance inflation due to collinearity with broader scope, providing assessment for both individual parameters and contrasts of parameters. Using this measure we introduce and briefly assess common modeling compromises, relating these compromises to our own and ABCD’s modeling choices for the MID task. Notably, the modeling approach used in the ABCD study is not unique, as other research groups have employed comparable models in other MID studies.

Collinearity can be addressed during study design by evaluating efficiency and variance inflation factors (VIFs). For a contrast *c* of parameter estimates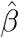, the variance is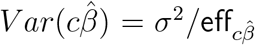, where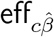is contrast efficiency (Appendix A) and *σ*^2^ is residual variance. This direct link means doubling efficiency halves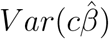, boosting power. However, efficiency only ranks task designs and offers no absolute benchmark for evaluating a single model.

In a linear model, *Y* = *β*_0_ + *β*_1_*X*_1_ + … + *β*_*p*_*X*_*p*_ + *ϵ*, the traditional VIF (tVIF) for 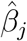 is the ratio of 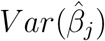 in the full model to that in the reduced model, *Y* = *β*_0_ + *β*_*j*_*X*_*j*_ + *ϵ* (derivation in Appendix A). This measures variance inflation due to *regressor* correlations, with benchmarks typically set at 5- or 10-fold increases (James et al., 2013, Section 3.3). However, tVIF applies only to individual parameters and can miss collinearity arising from overlapping condition signals when certain model parameterizations are used. We introduce a *contrast*-based VIF (cVIF) that captures this overlap and estimates VIFs for parameter contrasts (Appendix C). Further motivation for cVIF appears below and in Appendix B.

Figure 1 uses cVIF to illustrate common challenges in defining fMRI task regressors. Each panel presents a modeling question with two design options, the top representing a typical starting point, with cVIF values shown in the figure titles. For clarity, each design image shows single event presentations, but cVIFs are computed based on the number of trials that fit within a 50s period. While an improved task design can avoid many issues, these examples assume the data are already collected and the task design cannot be changed, requiring the researcher to make trade-offs to best use the available data. The simplified examples here are meant to exemplify challenges faced in our own and ABCD’s modeling of the MID task.

**Figure 1:**
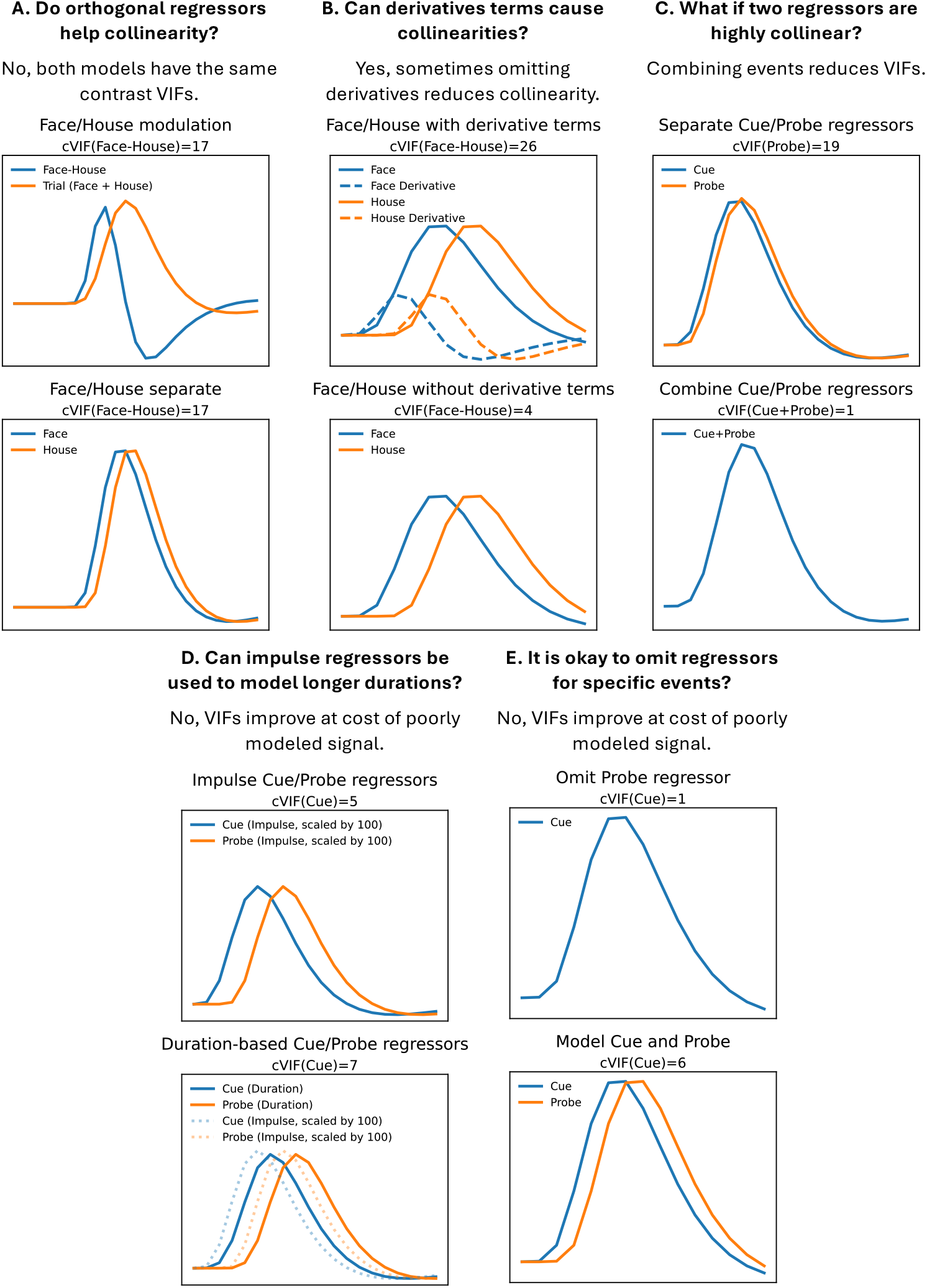
Different modeling strategies used when collinearity is encountered. Each panel represents a different scenario where the top model is a common starting point and the bottom model is an equivalent (Panel A) or improved compromise in model setup (Panels B-E). Only a single trial is shown for each event for the purposes of illustrating the collinearity, but the cVIFs are based on a 50s long scanning run. The cVIF for the contrast of interest is displayed at the top of each panel and is based on the algorithm presented in Appendix C.

Panel A of Figure 1 demonstrates the motivation for cVIF using two statistically equivalent models.

The top model uses orthogonal regressors with an ideal tVIF of 1, yet shows substantial contrast variance due to condition signal overlap (cVIF = 17). Details are provided in Appendix B. The reparameterization strategy to reduce regressor correlations is not used in ABCD’s MID model, but it is common in other MID studies and task fMRI in general. Re-parameterizing does not reduce the true variance inflation, as cVIF reveals, and relying on tVIF alone may give a misleading impression of improved model quality. Caution is needed when model parameterizations are described as remediating collinearity.

Panel B of Figure 1 shows a case where omitting derivative regressors is a pragmatic choice. Although not a standard practice, modeling a condition using a canonical HRF and HRF derivative regressor pairs can capture signals with shifted onsets. When conditions are brief and closely spaced, a shifted version of one condition’s regressor can overlap with another, which can drive collinearity when derivative regressors are added. A nonzero parameter estimate for a derivative term implies that the modeled response for a condition should be shifted in time. Although it is not common to use the derivative parameter to adjust amplitude estimates, as shown in Calhoun et al. (2004), including a derivative regressor still adjusts the estimates of other conditions for any increased overlap caused by the shift. In this example, cVIF drops from an unacceptable 26 to 4 when the derivatives are omitted, making omission preferable to discarding the data. If derivatives are omitted when thought to be necessary, the reasoning should be clearly stated. Alternative HRF modeling strategies using basis sets (Lindquist & Wager, 2006; Zarahn, 2000) are not covered here but can also suffer from collinearity due to condition overlap. As described in the Methods section, for our model, derivative terms were excluded due to high cVIFs, some over 100. In the Discussion we focus on the limitations introduced due to this decision, which impacts the flexibility to study certain contrasts using any statistical model due to constraints imposed by the study paradigm.

Panel C of Figure 1 shows a commonly occurring problem in many fMRI tasks where Cue and Probe regressors are highly collinear, preventing their separate analysis. A pragmatic, though im-perfect, solution is to merge them into a single event from cue onset to probe offset. This resolves collinearity but sacrifices interpretability, as the cause of activation cannot be disentangled.

A collinearity reduction strategy to avoid is modeling a condition with an impulse regressor instead of the true neuronal signal duration (Panel D, Figure 1). Mismodeling neuronal signal duration can shift the regressor peaks (shown in bottom model, Panel D, Figure 1), biasing the amplitude estimate of that condition and mismodeling the signal, which can bias other condition estimates. Our simulation analysis will show how the impulse regressors used in the ABCD model can bias contrast estimates. There may be limitations in knowing the true neural signal duration and it may vary across the brain (Summerfield et al., 2006), but the model should reflect the best current knowledge of the task. If the true activation duration is thought to be less than the event duration, then it is still likely necessary to model the remaining event duration’s period in a secondary regressor or design the task so the duration of the event aligns with expected duration of the neural signal.

Another important aspect of modeling neural duration is that duration and amplitude are confounded in the convolved BOLD response: for durations under 2s under conditions of equal intensity of neural activity, a neural event twice as long yields roughly twice the BOLD amplitude. For example, if response times differ between conditions and are not accounted for in the model, condition differences may be misattributed to increased neural activity rather than duration. This can be addressed with a single RT regressor (Mumford et al., 2024). In this MID data, both RT and duration differences in the Fixation and Probe events may confound trial outcomes (hit/miss), which will be discussed further below. None of these were modeled in the ABCD model.

Omitting regressors to address collinearity (Panel E, Figure 1) is a problematic practice due to its potential to significantly bias estimates. While the impact of unmodeled confounds in linear regression is well established (Greene, 2003), our simulations further demonstrate that this can be a substantial source of bias. More broadly, omitting known variables violates the assumption of “exogeneity” in linear models, which states that the error term in the model should be uncorrelated with any independent variable (whether or not it is included in the model). If this fails, as it can when an omitted variable is correlated with both an included independent variable and the dependent variable, then the statistical estimates can be invalid (Antonakis et al., 2010). The ABCD model omits some events and our simulations establish the potential for bias in contrast estimates.

Next, we review the history, design and traditional modeling decisions of the MID task. We do not discuss the mixed selectivity of brain regions for distinct components of the task, which are outside the scope of this paper (Bartra et al., 2013; Oldham et al., 2018).

### 1.2 The history and design of the MID task

The MID task was inspired by preclinical work that suggested dissociable affective responses to “reward anticipation”, “reward feedback” and error-related “learning” (Knutson & Wimmer, 2007; Schultz, 1998). Knutson’s original task design focused on mapping these distinct stages of reward-guided behavior to their neural correlates in humans across the distinct temporal events in the task: anticipation and feedback (Knutson et al., 2001). Knutson and colleagues reported that the reward anticipation stage was associated with heightened activity within the ventral striatum with a particular emphasis on the Nucleus Accumbens (NAc), while reward feedback was associated with heightened activity of the ventromedial prefrontal cortex (vmPFC; e.g., Knutson et al. (2003)). Researchers administer the task due to its postulated ability to distinguish between distinct reward valence systems, positive to *approach* and negative to *avoid* (Knutson et al., 2014). As a result, the MID has been adopted by many groups as a robust measure of incentive motivation processing (Bjork, 2020, p. 422) and included in the ABCD Study based on its ability to delineate processes of anticipation and receipt of reward (Casey et al., 2018, pp. 46-48).

The trial-by-trial structure across MID task administrations is comparatively similar to the version described in Knutson et al. (2001) and the ABCD study (Casey et al., 2018). On each trial, participants can *win* or *avoid losing* a large or small amount of money, or no money is at stake (i.e., neutral condition). A cue (henceforth, Cue) first notifies the participant of the trial type (Win, Don’t Lose or Neutral) and the magnitude of potential gain or loss (e.g., $5/$0.20 in the ABCD study). Following Cue presentation for a fixed amount of time (2000 ms in the ABCD study), a fixation point (henceforth, Fixation) then appears for a fixed (Yau et al., 2012) or variable amount of time (from 1500 to 4000 ms in the ABCD study). Next, a probe (henceforth, Probe) appears briefly (150 to 500 ms in the ABCD study), requiring a rapid button response. A successful response during the Probe presentation results in winning or avoiding a loss. The Probe duration is adaptively modified to achieve a target accuracy, with the ABCD study targeting 60%, accuracy and adjusting the duration every third trial based on the accuracy of the last six trials across *all* Cue conditions (Casey et al., 2018; Chaarani et al., 2021). In other versions, the duration for Probe may be adjusted within trials of each Cue condition (Wu et al., 2014). Following the Probe, the participant receives feedback (henceforth, Feedback) indicating whether they were successful/unsuccessful (too fast/too slow) in responding during the allotted Probe window and the resulting reward for that trial using a fixed (Knutson et al., 2003) or variable duration (Casey et al., 2018) (in the ABCD study, 1950 ms minus that current trial’s Probe duration).

### 1.3 *Variability* in fMRI time series models of the MID task

Each trial during the MID task aims to temporally separate distinct components of reward processing (Knutson & Wimmer, 2007). The proposed theoretical framework is that the anticipatory phase serves as the prediction for the anticipated reward (or expected value) and the outcome serves as the Feedback or prediction error.

While the Cue and/or Fixation components of the task are commonly modeled as “anticipatory”, the definition of this construct remains variable in the MID task literature (M. I. Demidenko et al., 2024). Whereas the Cue specifies the reward type and amount for a given trial, the Fixation may subserve a preparatory signal for the behavioral response during the Probe. As a result, some have referred to the Cue and Fixation as the anticipation of prospect, or the *motoric anticipation* (Andrews et al., 2011; Balodis et al., 2012; Garrison et al., 2017), separating it from the *anticipation of incentive* which occurs during the phase after the Probe and before the Feedback in versions of the task where a jitter is included between the two components. If the Cue serves as the expected incentive of a given trial and the Fixation serves as a preparatory response for that value, these adjacent components may contain overlapping but distinct cognitive processes. As a result, there is variability in how time series models of the MID task are specified, with differing definitions of anticipation and different events being left unmodeled.

Typically, the time series model of the MID task only includes regressors for the anticipation and Feedback components (Büchel et al., 2017; Srirangarajan et al., 2021). However, in some studies, the subject-level models omit modeling the Feedback component entirely (Weiland et al., 2013; Yau et al., 2012). The Probe is frequently omitted from the time series model, with some exceptions (Beck et al., 2009; Bjork et al., 2011; Knutson et al., 2005). Anticipation has been defined variably, including: 1) start and duration of the trial’s Fixation events, leaving Cue unmodeled (Knutson et al., 2003); 2) the start and duration of the trial’s Cue events, leaving Fixation unmodeled (Joseph et al., 2016); 3) an impulse response at the start of the trial’s Cue events, leaving Fixation unmodeled (Hagler et al., 2019); or 4) the start of the trial’s Cue events and *combined* duration of the trial’s Cue and Fixation events (Balodis et al., 2012; Martz et al., 2016). In some cases, like in the ABCD study (Hagler et al., 2019), researchers have included temporal derivatives for each of the modeled events (Joseph et al., 2016). It is rare for response times to be modeled in the MID task, although RT adjustments to the anticipatory signal were made in Knutson et al. (2005). This variability in designs (Wilson et al., 2018), definitions and model specifications makes it challenging to compare the model-derived estimates, which can vary substantially in their magnitude and reliability across the different specifications (M. I. Demidenko et al., 2024).

Another variable modeling strategy involves the definition of Miss trials, which is rarely documented in manuscripts, so we have relied on personal communications with other MID researchers for clarification. Hit trials are consistently defined as a button press during the Probe phase, but the other two outcomes—Too Soon (a button press before the Probe phase) and Too Slow (a button press absent or after the Probe offset)—are used inconsistently. Some researchers classify both Too Soon and Too Slow as Miss trials, while others define Miss trials as Too Slow only, discarding Too Soon trials or modeling them as a separate regressor of no interest (Büchel et al., 2017). Leaving these unmodeled has the potential for them to bias estimates of the subsequent feedback component.

### 2 Methods

A subset of the ABCD dataset was used for both simulated and real data analyses. Task timings and participants’ behavioral responses were used to generate design matrices and simulate fMRI data, which were analyzed using two modeling strategies. These models were then applied to the real data, allowing for a comparison of group activation maps from commonly used MID contrasts.

### 2.1 ABCD Data

#### Participants

The ABCD sample used here is from the raw ABCD-BIDS Community Collection (ABCC, Feczko et al. (2021)), which was accessed via the Minnesota Super-computing Institute and Amazon Web Services. The ABCD study protocol acquired structural and functional volumes across 21-sites and three separate scanner vendors (Casey et al., 2018). The focus here is on the 2-year follow-up data for two reasons. First, earlier versions of the MID E-prime procedure administered during baseline did not record reaction times (RTs) for miss trials. Second, the sample used here is a sub-sample assessed for quality control in a previous analysis (M. I. Demidenko et al., 2024) including successful field map correction, co-registration between anatomical and functional data, mean framewise displacement < .9, and availability of e-prime data and adequate task performance. For the final analytic sample (N = 500), the average age of the participants is 12.03 years-old (SD: 0.66) and 47% are Female (N = 235). All participants’ data are from SIEMENS scanners across 13 different ABCD sites.

#### Monetary Incentive Delay task

These analyses focus on the modified version of the MID task (Knutson et al., 2001) that is administered in the ABCD study protocol (Casey et al., 2018). A trial of the MID task includes Cue (1781-2039ms), Fixation (1500-3666ms), Probe (172-497ms) and Feedback (1473-1789ms) components (These ranges deviate slightly from the values reported in Casey et al., 2018 for reasons described below). There are five Cue types: Win $5 (Large Win), Win $0.20 (Small Win), No Money at Stake (Neutral), Don’t Lose $5 (Large Loss), and Don’t Lose $0.20 (Small Loss). Based on the subject responses, Feedback is classified as either a Hit (successful probe window response) or Miss (Too Soon/Too Slow response). Each run consists of 50 sequential trials (10 per condition) that are presented in pseudo-random order. The ABCD study protocol did not include inter-trial intervals.

During the development of these analyses we identified issues with the ABCD acquisition and timing files. First, since the .*Duration* columns in the E-prime files reflect the requested rather than the true event screen-display durations, we use the .*OnsetToOnsetTime* column to determine the true event duration (Psychology Software Tools, 2024). In the case of the Feedback events there isn’t a .*OnsetToOnsetTime* column so we use the difference between the onset of the next trial’s Cue and the onset of the current trial’s Feedback as the event duration, which is consistent with the .*OnsetToOnsetTime* definition (Psychology Software Tools, 2024). Second, the original response time estimates within the Eprime files, denoted “OverallRT” were based on flawed event duration values for Miss trials, so we recomputed them using updated trial durations. This adjustment affected only the Too Soon and Too Slow response times, when present. A trial is classified as Too Soon if any button press occurs before the Probe. The recording of response times on Too Soon trials by the E-prime script depends on the timing and number of button presses. If only a single button press occurs on a Too Soon trial, no response time is recorded. However, if additional button presses occur after the Probe onset, the response time of the first press is recorded for that trial. Lastly, as an aside, trial onset timing values for a subset of ABCD data were misspecified due to an incorrect estimation of calibration volumes in all GE data; see Supplemental Section S1 for a more detailed description of this issue. This issue impacts all ABCD and ABCC GE data released prior to May 2023 and were used to generate the statistical models in those releases. The issue is expected to be fixed in the ABCD Release 6.0 and future ABCC releases.

The time series models studied are defined in terms of either impulse functions (i.e., Dirac delta) or boxcar functions convolved with a double gamma hemodynamic response function (HRF), specifically the `spm_hrf` function from Nilearn (Abraham et al., 2014). In the ABCD study, a Miss is defined as either Too Slow or Too Soon. We follow that definition in our simulations to reflect the modeling strategy used to date in the ABCD study (Hagler et al., 2019). Here, we compare two models:

- **CueFeedback model** (Hagler et al., 2019). This parallels the model used by ABCD. All regressors are based on impulse functions at event onsets. There are a total of 5 Cue regressors (large loss, small loss, large win, small win and neutral) and 10 Feedback regressors (each cue type split by hit/miss). Each of these 15 regressors is paired with its temporal derivative for a total of 30 task-related regressors (left panel, Supplemental Figure S2).
- **Saturated model**. In our proposed model, regressors are based on boxcar functions using the event onsets and event durations. There are 5 Cue regressors split by cue type (duration: 1781-2039 ms), 5 Fixation regressors split by cue type (1500-3666 ms), 3 Probe regressors split by win/loss/neutral (172-497 ms), 1 response time (RT) regressor (uses Probe onset and the RT as the duration, between 105-1581 ms) and 10 Feedback regressors split by cue type and hit/miss (1473-1789 ms) (right panel, Supplemental Figure S2).

For the CueFeedback model, the use of impulse regressors reflects the issue described in Figure 1D. Also, the CueFeedback model omits regressors for Fixation, Probe and RT, which reflects the issues presented in Figure 1E. The impact of these decisions will be studied in the simulation analysis

For the Saturated model we did not include derivative terms as their addition led to 30 of the 49 model parameters having excessive cVIFs (> 20). For example, commonly studied contrasts comparing large win to neutral and large loss to neutral for Cue trials, have cVIFS > 200. This tradeoff reflects the dilemma illustrated in Figure 1B: either abandon the analysis or omit derivatives, recognizing that the model will not account for signal shifts overlapping with subsequent events. Although there is evidence that a more flexible model (including derivative and dispersion regressors) is justified, the MID task’s design makes this difficult to implement in practice due to collinearity. We return to this limitation in the Discussion, where we offer recommendations.

An illustration of the two models and the estimated components are presented in Figure 2. Panels A and B illustrate regressors for a single trial within each design where blue, yellow, purple, red and green correspond to Cue, Fixation, Probe, Feedback and RT, respectively. The curves are the regressors, using a dashed line for derivative regressors. The color blocks along the x-axes indicate the onset and durations that were modeled for each event. Panel C provides examples of brain maps that can be estimated for each of the model types.

**Figure 2:**
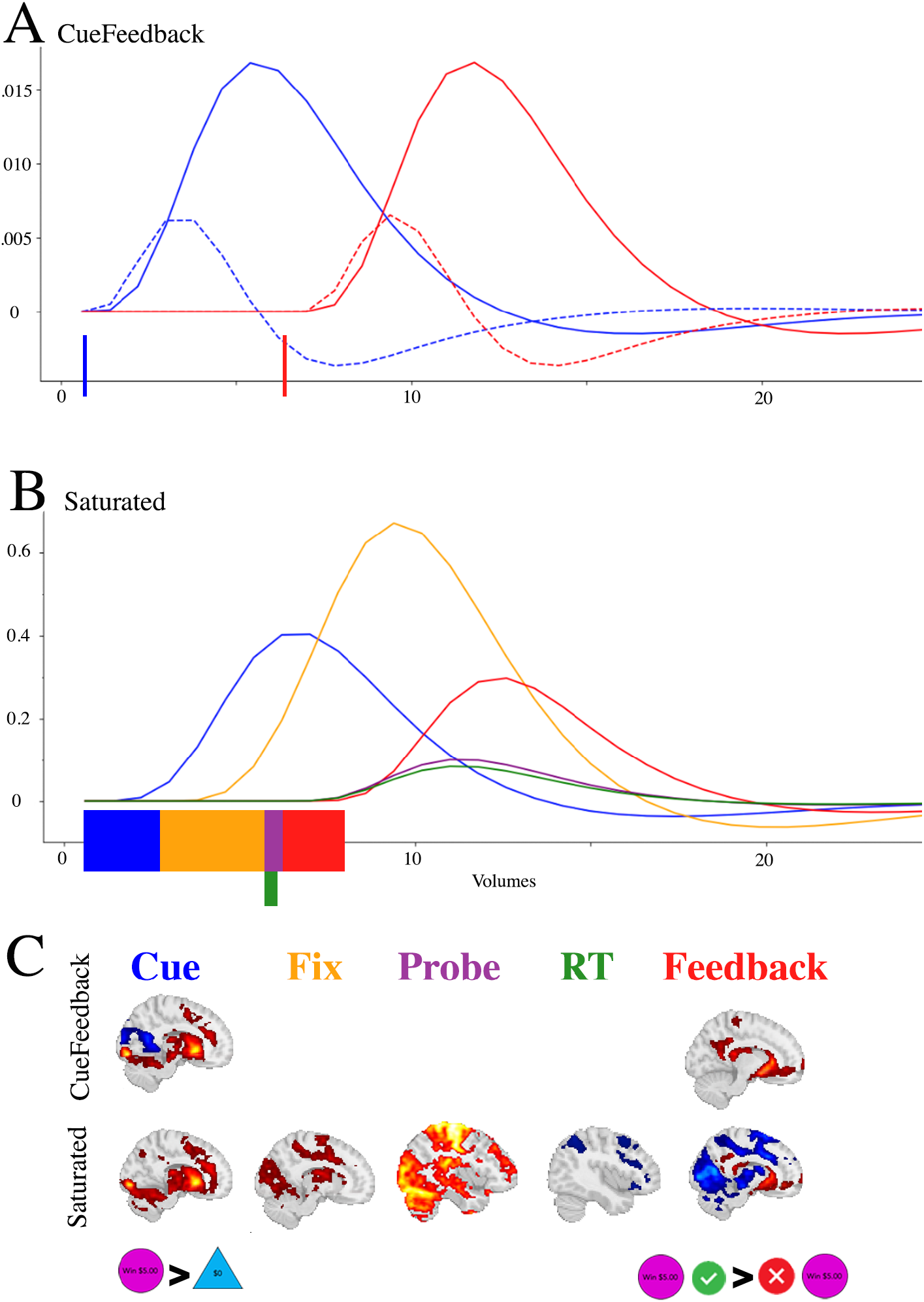
Illustration of the onset times, durations and estimated BOLD response for task components in the (A) CueFeedback and (B) Saturated model; and the (C) estimated maps between the CueFeedback and Saturated models. The colored blocks along the x-axes in Panels A and B represent the onset and durations being modeled and the dashed lines in Panel A correspond to the derivative regressors. The example activation maps are subset from Supplemental Figures S11, S13 and S14. The dashed lines in Panel A correspond to the temporal derivative (see Supplemental Figure S2). Note: in this example the Probe is for a Win condition.

### 2.2 Simulated data analysis

Code for all simulations is publicly available (R. Poldrack & Mumford, 2025). Simulated data represented time series across multiple subjects within a single voxel. Design matrices were constructed by concatenating each subject’s two runs into a single run. Regressors were constructed by convolving boxcar (or delta functions) with the spm HRF. Synthetic data were generated by multiplying the Saturated model design matrix by the desired true signal components, and then adding independent Gaussian noise to the true signal. The time resolution matched that of the real data (800 ms). Independent noise is not a plausible assumption for fMRI data, but it does not impact the overall simulation results and allows for the use of a simple ordinary least squares model here. Between-participant variability was introduced by sampling the true signal components from a Gaussian distribution with the intended mean (Table 1); the standard deviation of the between-participant and within-participant variability were set as equal to 1.5 and 1, respectively. If the fMRI time series model is defined by

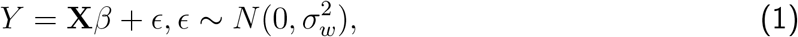

where *X* is the design matrix, *β* is the vector of model parameter and *ϵ* is the error term with a variance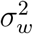, the within-subject variance of a contrast of parameter estimates, **c***β*, is given by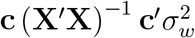. If the between-subject variance is defined as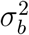, then the total mixed effects variance for a first-level contrast of parameter estimates is

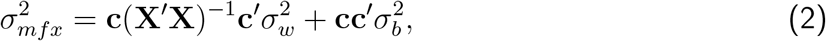

and the ratio of the square root of the total variance to the square root of the within-subject variance is given by:

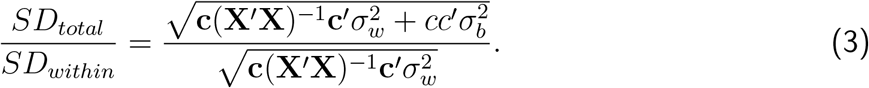

**Table 1:** Simulation settings. Each row corresponds to a different simulated data set, where the nonzero parameters are specified in the second column.

| Condition(s) of focus | Nonzero parameters |
| --- | --- |
| Null model | (all betas 0) |
| Cue (Win) | Cue LargeWin = 0.22,<br>Cue SmallWin = 0.22 |
| Fixation (Win) | Fixation LargeWin = 0.22,<br>Fixation SmallWin = 0.22 |
| Probe (Win) | Probe Win = 0.85 |
| Response Time | Probe: RT = 0.35 |
| Feedback | Feedback LargeWinHit = 0.25,<br>Feedback LargeWinMiss = 0.25 |

Our chosen within- and between-participant variances yielded values of 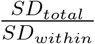 ranging between 2-4.2 across the condition difference contrasts studied and design matrices across subjects, consistent with what we have done in other published simulations (Mumford et al., 2024). For each simulated data set, aside from the null data simulation, only a subset of parameters had nonzero values as shown in Table 1. Parameter values were chosen manually such that the power of detection, for that parameter, was approximately 80% for the group average. The contrasts we studied in the simulation analysis, which were also the main focus of the real data analysis, are shown in Table 2. To avoid redundancy, we focused on win-based contrasts, omitting their loss-based counterparts (e.g., included Cue:LW-Neut^*^ but omitted Cue:LL-Neut^*^), as simulations introducing signal for Win trials yield results similar to those for Loss trials. Note the contrasts names with asterisks indicate the contrasts of focus for the ABCD study. For each parameter setting, a total of 1000 data sets containing 500 subjects each was simulated and analyzed. Error rate and bias were estimated for contrasts that were null for that simulation setting.

**Table 2:** Contrasts used in simulation analyses and of primary focus in the real data analyses and contrasts used.

| Contrast Name | Contrast Definition |
| --- | --- |
| **Cue: LW-Neut\*** | $1 \times \text{Cue LargeWin}$<br>$- 1 \times \text{Cue Neutral}$ |
| **Cue: LL-Neut\*** | $1 \times \text{Cue LargeLoss}$<br>$- 1 \times \text{Cue Neutral}$ |
| **Cue: LW-Base** | $1 \times \text{Cue LargeWin}$ |
| **FB: WHit-WMiss\*** | $.5 \times \text{Feedback LargeWinHit}$<br>$.5 \times \text{Feedback SmallWinHit}$<br>$- .5 \times \text{Feedback LargeWinMiss}$<br>$-.5 \times \text{Feedback SmallWinMiss}$ |
| **FB: LHit-LMiss\*** | $.5 \times \text{Feedback LargeLossHit}$<br>$.5 \times \text{Feedback SmallLossHit}$<br>$- .5 \times \text{Feedback LargeLossMiss}$<br>$- .5 \times \text{Feedback SmallLossMiss}$ |
| **FB: LWHit-NeutHit** | $1 \times \text{Feedback LargeWinHit}$<br>$- 1 \times \text{FeedbackNeutralHit}$ |
| **FB: LWHit-Base** | $1 \times \text{Feedback LargeWinHit}$ |

\* Contrast distributed by ABCD

### 2.3 Real data analysis

#### Preprocessing

The data were preprocessed using fMRIPrep v23.1.4 (Esteban et al., 2022). In short, a T1-weighted image was corrected for intensity normality, skull-striped via Nipype’s ants-BrainExtraction.sh workflow and spatially normalized to MNI152NLin2009cAsym standard space using non linear registration via AntsRegistration. For each of the two MID BOLD functional runs, the following sequence of steps were performed. A reference volume and its skull-stripped version were generated, a fieldmap was estimated and aligned with rigid-registration to the target EPI (echo-planar imaging) reference run and each BOLD run was corrected for signal distortion using top-up. The BOLD reference image was co-registered to the T1w image using bbregister using boundary-based registration (Greve & Fischl, 2009) with six degrees of freedom. The BOLD time-series were resampled into standard space, resulting in a preprocessed BOLD run in MNI152Nlin2009cAsym space. Head-motion parameters (transformation matrices and six corresponding rotation and translation parameters) were estimated before any spatiotemporal filtering using mcflirt (Jenkinson et al., 2002) and expanded with the inclusion of temporal derivatives and quadratic terms for each (Satterthwaite et al., 2013). The time-series data were filtered using a discrete cosine filter with 128s highpass cut-off. For each BOLD run, an associated confound file was generated with the estimated motion parameters and the cosine basis set that were used in subsequent subject-specific GLMs.

#### Model estimation

Code used for the model estimation is available in M. Demidenko et al. (2025). Within-run and within-subject models were estimated using *Nilearn* 0.9.2 (Abraham et al., 2014) in Python 3.9.7. Inference for group-level 1-sample t-tests were carried out using cluster-based family-wise error rate (FWER) corrected statistics estimated with a permutation test accounting for the grouping of data within-site.

For each of the two runs, the within-run model fits a GLM to the subject-specific time series data using the function `FirstLevelModel` in *Nilearn*. In addition to the task regressors described above for the two models, cosine regressors calculated by *fMRIPrep*, corresponding to a 128s highpass filter cut-off, and the x, y, z translations, rotations and their derivatives (12 motion parameters), were included in the subject-specific design matrix. The data were spatially smoothed using a 5 mm FWHM smoothing kernel and the model included an AR(1) model of temporal autocorrelation. For each contrast of interest, *Nilearn*’s `compute contrast` was used to obtain a contrast estimate (`effect size`) and its estimated variance (`effect variance`). The within-subject model estimates average contrast estimates across the two runs. For within-subject estimates we used *Nilearn*’s `glm.compute fixed effects` with the precision-weighted option to obtain a weighted-average of the contrast estimates across runs.

For the group-level 1-sample, 2-sided t-test, we employed a cluster-size based permutation test to control for family-wise error rate (FWER) while accounting for site effects. The procedure generates a null distribution by performing sign-flipping permutations stratified by scanner site (yielding 2^13^ = 8192 possible permutations), which preserves the multi-site structure of the data (Winkler et al., 2015). Clusters were defined using 6-connectivity, face-adjacent voxels, in 3D space. For each permutation, we recorded the maximum cluster size after applying a cluster-forming threshold of *t* = 3.1 (two-sided). The 95th percentile of this null distribution of maximum cluster sizes was then used to determine significant clusters in the observed data, ensuring FWER control at *p* < 0.05. This approach provides robust statistical inference while appropriately accounting for the non-independence introduced by the multi-site acquisition. For display purposes the *t*-statistics were converted to Cohen’s *d* estimates using:

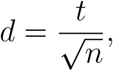

where *n* is the number of subjects. Cohen’s *d* values are displayed for voxels in significant clusters. Unthresholded Cohen’s D maps are in the main manuscript and thresholded images can be found in the Supplementary Materials.

#### Region of Interest (ROI) Estimates

ROI and model-specific contrast estimates were extracted for the Left and Right NAc. The Left and Right NAc are from the thresholded Harvard-Oxford subcortical atlas (*nilearn* Harvard-Oxford: sub-maxprob-thr50-2mm) using the procedure described in Supplemental Section S6. To extract ROI estimates from the statistical maps, for each contrast the voxels were masked and averaged within the Left and Right NAc.

## 3 Results

### 3.1 Simulation Study

Supplemental Figure S3 displays the cVIFs for regressors and contrasts using our new cVIF algorithm for contrasts (Appendix C). Unexpectedly, even without an ITI in the ABCD version of the task, the Saturated model with regressors for all MID trial events does not exhibit problematic collinearity for contrasts of interest (indicated by asterisks: Cue:LL-Neut*, Cue:LW-Neut*, FB:LHit-LMiss*, FB:WHit-WMiss*). Although the cVIFs are above 15 for the Win/Loss/Neutral Probe regressors, these activation estimates are not studied in the MID task literature. Also, as will be seen in our real data analyses, the signal for these contrasts is very large, outweighing any concerns of variance inflation, since increased variance is more problematic in estimates with low signal to noise ratio. For the contrasts of interest (Table 2), only the cVIF for FB:LWHit-Base is slightly elevated (9.66). However, this contrast is not commonly studied in the MID task literature and is used here for illustration purposes only. While elevated VIFs can suggest reduced statistical power, the inferences drawn from the analysis remain statistically valid, albeit at an increased risk of outliers. When VIFs exceed the commonly accepted threshold of 5 for a contrast of interest, we recommend the researcher carefully inspect the data for outliers, for example by voxelwise analyses of whether each subject’s estimate is 2 or 3 standard deviations from the mean (across subjects) and checking whether a subject has outlying estimates across a large portion of voxels. Python code for carrying out this evaluation can be found at github.com/jmumford/fmri-outlier-detector. Carpet plots of contrast estimates across subjects can also be inspected for outliers. Identified outliers should be carefully monitored if included in further analyses using proper outlier influence assessment tools for the analysis used (e.g., Cook’s D for correlations between brain and behavior measures (Neter et al., 1996)).

Efficiency differences between the two models are shown in Supplemental Figure S4. Aside from the FB:LWHit-Base and FB:LWHit-NeuHit contrasts, the efficiencies between the two models for contrasts of interest are similar. Notably, the commonly studied contrasts, FB:LHit-LMiss^*^ and FB:WHit-WMiss^*^, have efficiencies within 6%. Given that between-session and between-subject variances typically dominate overall variance in group-level models, these small efficiency differences are unlikely to affect group-level results. As such, for these two contrasts it is likely a good assumption that activation differences between the models are primarily driven by bias differences rather than power differences.

Recall that the CueFeedback model excludes regressors for Fixation, Probe, and RTs, leaving unmodeled variability that can bias other model parameters. Additionally, if these unmodeled events vary in neuronal activation durations across trial and outcome types, this can unevenly bias parameters, as duration differences in the neuronal signal may appear as amplitude differences in the BOLD signal (Mumford et al., 2024). Figure 3 shows that Probe, Fixation, and RT durations differ between trial outcomes (Hit/Miss), indicating a potential for biases on contrasts between Hit and Miss. To clarify details about the RTs, the measured RTs are button presses after Probe onset, with Too Soon RTs indicating a secondary press within a trial and Too Slow RTs corresponding to presses after Probe offset. On average, Probe durations are 26-33ms longer for Hit trials, Fixation durations are 63-160ms longer for Hit trials and RTs are 78–100ms longer for Miss trials.

**Figure 3:**
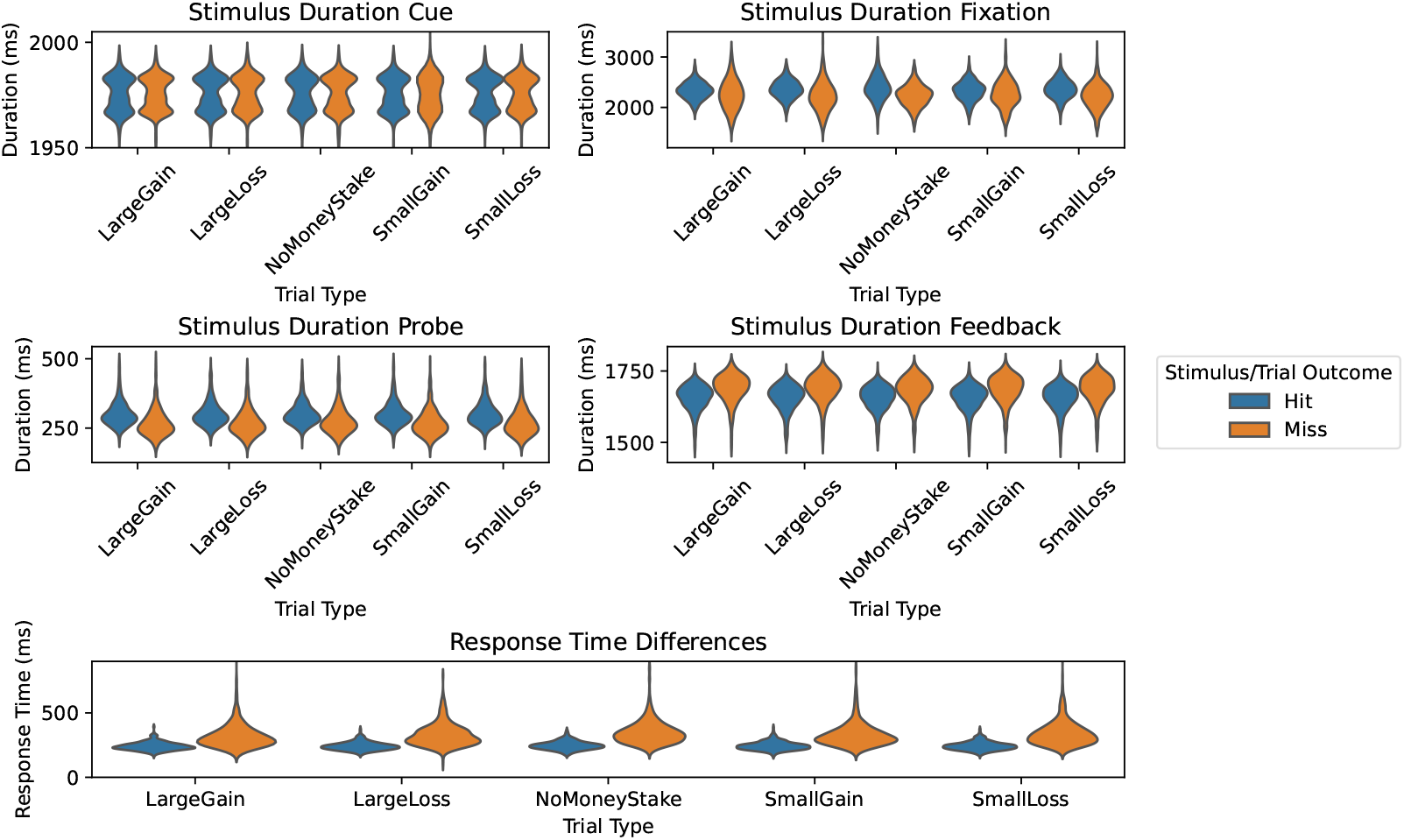
Event durations and response times split by trial type and trial outcome. Differences in duration or RTs between trial outcomes indicate the potential for contrasts comparing hits and misses to be biased.

Supplemental Figure S5 further breaks down Miss trials into Too Soon and Too Slow, highlighting notable differences. Compared to Too Slow trials, Fixation durations are much longer (700ms) for Too Soon trials, Probe durations are slightly longer (10ms) for Too Soon trials and RTs are shorter (42ms) for Too Soon trials. On average, trials are 57% Correct, 36% Too Slow, and 7% Too Soon. Given these differences as well as differences in the cognitive processes that are occurring during Too Slow and Too Soon, a more specific model would separate Feedback regressors into Correct, Too Soon, and Too Slow, with a focus on comparisons between Correct and Too Slow while Feedback for Too Soon would be treated as a nuisance regressor. We have not explored such a model, for the sake of brevity, but we suggest that it be considered in future MID research.

Figure 4 illustrates how biases can be introduced into contrast estimates when using the Cue-Feedback model. Recall, contrasts with asterisks are contrasts distributed by the ABCD group, while contrasts without asterisks have been studied elsewhere but were not included in ABCD data releases. As expected, there are controlled biases for the Null setting for both models and the Saturated model has controlled biases for all simulated settings. When signal is introduced for the Cue events during Win trials (second row, Figure 4), significant positive bias is present for FB:LWHit-Base in the CueFeedback model, indicating the impulse regressor’s inability to capture all of the Cue-based activity. Additionally, bias in the CueFeedback model is also present for FB:LWHit-NeutHit, although it is not significant. As expected, since there are no Fixation regressors in the CueFeedback model, when Fixation signal is introduced for Win trials (third row, Figure 4) the unmodeled neural signal biases the CUE:LW-Neut^*\**^, FB:LWHit-Base and FB:LWHit-NeutHit. Introducing signal for Probe on Win trials (4th row) leads to significant bias in FB:LWHit-Base and FB:LWHit-NeutHit contrasts. Although it is not significant, a positive bias in FB:WHit-WMiss^*\**^ is present and is likely driven by the longer event durations for Hit Probe trials compared to Miss (Figure 3). Note that we consistently introduce signal associated with approximately 80% for each simulation, but the true Probe activation in the real data is quite large implying these biases would be larger in real data. Adding RT signal introduces a significant bias in the CueFeedback model’s estimate of FB:LWHit-Base. When Feedback signal is added, although the impulse regressor of the CueFeedback model may not capture all of the signal, the randomization of the trial order prevents biasing in the Cue-based contrasts.

**Figure 4:**
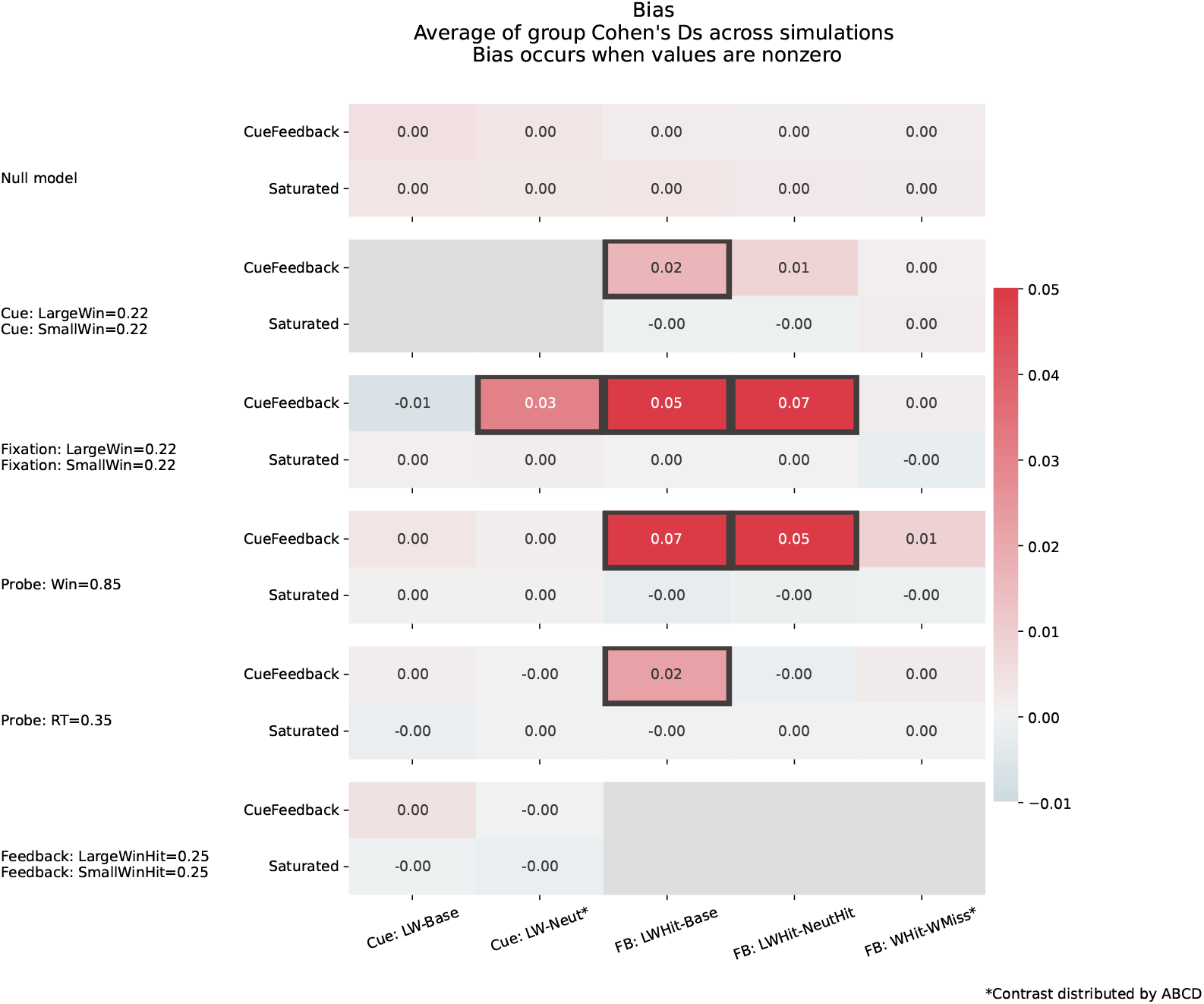
Bias for contrast estimates across simulation settings and models. Gray squares around cells indicate the Type I error rate was inflated, while the color reflects the average t-statistic, where a value of 0 reflects no biasing. Each row corresponds to a different model where the contrast estimate bias is shown for the CueFeedback (top) and Saturated (bottom) models. The solid gray cells indicate that a parameter involved in that contrast was nonzero and so error rate is not estimated.

#### 3.1.1 Real-world Data Analyses: Effect of model choice with the MID task

The within-run, within-subject and group-level analytic code, and the Jupyter notebook used to generate the associated tables and figures from the estimated maps are available in M. Demidenko et al. (2025). All within-run, within-subject and group maps use data from N = 500 ABCD sample participants. The resulting *t*-stat and Cohen’s *d* estimated images (corrected and uncorrected) across each contrast for the group and paired *t*-test between models are available on Neurovault: identifiers.org/neurovault.collection:20608.

##### Behavioral Results

Participant’s Probe response accuracy and response times across Cue conditions are reported in Supplemental Figure S10 and in Figure 3, respectively. As expected, Probe response accuracy differs across Cue events and mean RT values differ across Hit/Miss Feedback events.

##### Comparing Models: Whole-Brain Activation Results

The seven contrasts of focus for these real data analysis are defined in Table 2, while fourteen additional contrasts and their results are included in the Section S8 of the Supplemental Materials. The unthresholded activation maps for each model as well as their paired comparisons for these main seven contrasts are shown in Figure 5, the thresholded maps, using a cluster-based permutation accounting for site using 8192 permutations, are shown in Supplemental Figure S14 and the maps for the complete list of contrasts are presented in Supplemental Figure S11.

**Figure 5:**
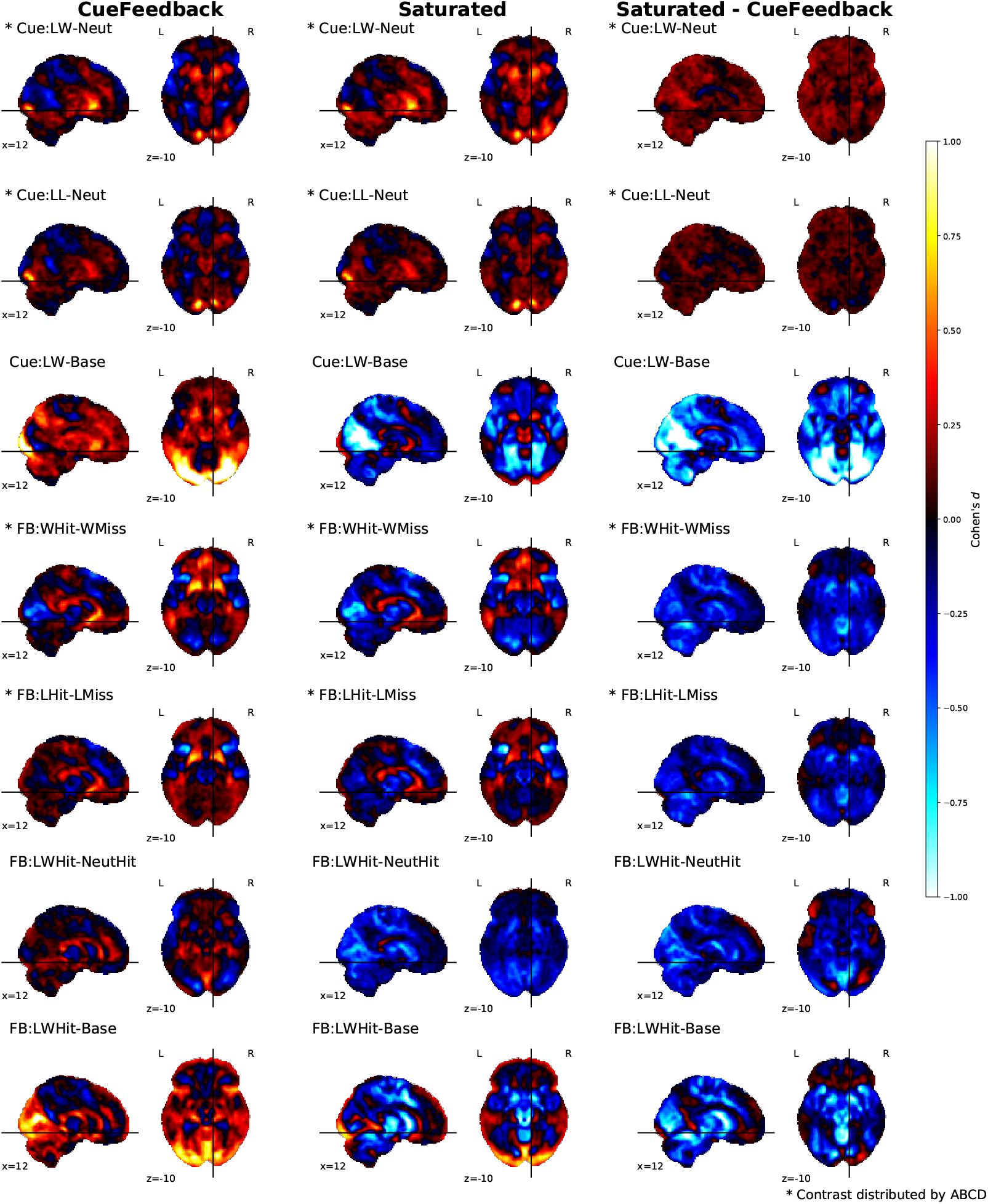
Cohen’s *d* [uncorrected] group-level activation maps across Cue (4) and Feedback (3) contrasts for the CueFeedback (shown on the left), Saturated (shown in the middle) and their paired difference (shown on the right). We note contrasts included with ABCD data releases Chaarani et al., 2021.

Across the Cue and Feedback contrasts for the CueFeedback and Saturated models, the group average activation maps are presented in the first two columns of Figures 5/S14. Consistent with the literature, the Cue contrasts (rows 1 - 3 in Figures 5/S14) demonstrate a broad activation of striatal, anterior insula, motor and visual areas. Likewise, consistent with the literature, the within condition Hit versus Miss contrasts (rows 4 & 5 in Figure S14) demonstrate activation in mPFC and striatal areas and, for the Saturated model, also meaningful deactivation in the anterior insula, visual and motor areas. The between-condition Hit contrast (row 6) demonstrates a broad deactivation in striatal, anterior insula, visual, anterior insula and motor areas in the Saturated model and positive activity in the rostral anterior cingulate cortex and lateral ventricles near the caudate nucleus in the CueFeedback model. Higher deactivation in the Saturated model is also present in the Feedback condition versus baseline contrasts. Which may point to misfit of the canonical HRF, a topic we return to in the Discussion section.

The CueFeedback and Saturated models meaningfully differed in the estimated BOLD activity at the group level across the seven contrast types. As is illustrated in the paired *t*-stat difference map in the right columns of Figures 5/S14, the CueFeedback and the Saturated model show meaningful differences in activation magnitude across contrast maps, particularly in the visual, anterior insula and striatal regions. During the Cue phase, the Cue:LW-Neut^*^, Cue:LL-Neut^*^ and Cue:L-Neut had meaningfully higher activity in the Left and/or Right NAc in the Saturated than CueFeedback model (see Supplemental Figure S11). This likely reflects improvement in capturing the signal when modeling the event durations. Furthermore, the simulation results demonstrate a positive bias in the FB:WHit-WMiss^*^ contrast when signal is added to the Probe component and left unmodeled. For FB:WHit-WMiss^*^ and FB:LHit-LMiss^*^ (rows 4 and 5 of Figures 5/S14), a reduction in subcortical activity can be observed when modeling for the Probe conditions in the Saturated model. Given the efficiencies for these contrasts were almost identical in these two models and that both positive and negative estimates shift in the negative direction in the Saturated model compared to the CueFeedback model, these differences likely illustrate an adjustment for Probe-based biases in the Saturated model. If the differences related to power differences, the estimates in the Saturated model would have diminished toward zero. Differences in the Feedback phase reached small to medium effect sizes in multiple regions. For example, the mean effect size (Cohen’s *d*, using a cluster based permutation accounting for site, with 8192 permutations) for the activation reduction within the NAc region of interest during Feedback ranged from *d* = -.43 to -.25 and *d* = -.41 to -.27 for Left and Right, respectively, across four of the within-condition Hit versus Miss contrasts, and *d* = -.60 to -.46 and *d* = -.66 to -.50 for Left and Right, respectively, across Large Win Hit > Neutral Hit and Large Win Hit > Implicit Baseline contrasts. The paired difference between estimated z-statistics across subjects between the CueFeedback and Saturated model are presented in the Supplemental Figure S9.

In Supplemental Figure 13, we present the Cue and Fixation contrast maps for the Saturated model. For the Fixation component, in the between-condition contrasts (e.g., Large Win*>*Neutral), there is heightened activity in the visual, motor, dorsal striatal and posterior brain regions compared to the contrasts for the Cue component. Furthermore, as is illustrated for the Saturated model in Supplemental Figure S13, the Large Win > Implicit Baseline contrast map is largely negative for Cue and positive for the Fixation component. This illustrates that the Cue and Fixation components differentially impact the estimate of Baseline activity and should be accounted for in subject-specific models.

Supplemental Figure S12 shows activation maps for the average Probe, RT effect, and the pairwise probe differences between the Win (Circle), Loss (Square) and Neutral (Triangle) probe events. The Probe is associated with very high positive BOLD activity across the motor system and other regions. Furthermore, there is a meaningful difference in activity between the Probe events. For example, activity is significantly higher in each of Loss and Win compared to Neutral. The Probe RT is associated with decreased BOLD activity in a broad set of regions including frontal and parietal regions. While present, the magnitude of the RT effect in this ABCD sample is relatively small., possibly due to the relatively small range of reaction times in this task.

## 4 Discussion

Given the weak signal in task fMRI, relative to the noise, researchers optimize task designs and subject-level models of the BOLD time series data to reduce collinearity between task regressors and increase power to detect small effects. Although the omission of task components or mismodeling the estimated duration of BOLD activity are common practices in task fMRI modeling, as they may reduce collinearity, they may also introduce bias into parameter estimates. Using ABCD’s MID task, we quantified the potential for bias to be present in contrast estimates when using the CueFeedback model, which reflects the model used by ABCD. These simulations affirm previously known biases in Feedback activation comparisons between Cue type and identified new biases of contrast estimates in MID task modeling. Even though there are multiple, adjacent components within a trial, with no resting fixation in between trials, the collinearity of the Saturated model for contrasts of interest (marked by an aterisk in Table 2) is within the acceptable range, as measured with our cVIF procedure. In real-world data, the Saturated model elicits BOLD activity in expected regions for typical Cue and Feedback contrasts, accounting for adjacent task components. The ABCD Consortium task fMRI workgroup are considering incorporating elements of the Saturated model for future releases, using cVIF to evaluate different candidate models. Notably, since response times were not collected for all subjects, this limits the ability of the ABCD Consortium task fMRI workgroup to add an RT adjustment. Although this was found to have limited bias on the 1-sample t-test evaluations performed here, it is unknown whether RT confound may impact individual differences analyses.

### 4.1 Activation map differences

Some differences between the performance of the CueFeedback and Saturated models are clear, but others remain uncertain due to the structure of the MID paradigm. A major improvement of the Saturated model is the inclusion of previously unmodeled events—Fixation, Probe, and RT—all of which vary in duration between Hit and Miss trials. Fixation and Probe are also expected to differ in evoked neural responses across Cue types. As discussed in previous work (Andrews et al., 2011; Balodis et al., 2012; Garrison et al., 2017), the Fixation may conceptually differ from the Cue component. A variety of factors are influencing the statistics, including the previously mentioned biases, challenges in interpreting parameter estimates, and potential differences in power. However, our main results do not appear to be affected by reduced power.

#### 4.1.1 Power versus bias

Power differences between the two models cannot be assessed, as the parameter estimates differ in meaning between tasks and, more importantly, potentially biased estimates cannot provide meaningful power assessments. In some cases the efficiencies between the two models are so close that it is unlikely that power differences could explain the activation differences (FB:WHit-WMiss^*^ and FB:LHit-LMiss^*^). The other two Feedback condition comparison contrasts are discussed in more detail below, but those differences are also not likely due to power. For Cue-based condition differences (Cue:LL-Neut^*^, Cue:LW-Neut^*^) the Saturated model is less efficient, but has larger estimates, implying there is not likely a power issue. Each category of contrasts is further discussed below.

#### 4.1.2 Cue comparisons with baseline

The biggest impact due to change in parameter interpretation can be seen in the contrasts that compare to baseline. In the CueFeedback model, the “baseline” activation consists of activation related to Fixation, Probe and RTs and does not reflect a resting baseline (e.g., time at the beginning or end of the run when they are not performing a task, since no such null periods are present in this version of the task). In the Saturated model, the baseline is the actual baseline and so the contrasts comparing a parameter to baseline have very different meanings between the two models. Although contrasts versus baseline in MID studies are rare, they do occur (Andrews et al., 2011; Patel et al., 2013) and are recommended by some (Balodis & Potenza, 2015). Compounding this problem across MID studies is inconsistency in how anticipation is defined and what unmodeled neural activations are contributing to baseline. Sometimes the Anticipatory signal is defined as occurring during the Cue (leaving Fixation unmodeled) while other times it is defined as occurring during the Fixation (leaving the Cue unmodeled). These two modeling styles will produce estimated anticipation versus baseline effects that are the negatives of each other (M. I. Demidenko et al., 2024). By using the Saturated model, the interpretation of all contrasts in the model will change because they are now adjusted for Fixation, Probe and RT effects.

### 4.1.3 Cue-based condition comparisons

As expected, the CueFeedback model elicits activity in the direction and magnitude that is consistent with previous contrast estimates by the ABCD Consortium task fMRI workgroup (Chaarani et al., 2021). For example, in the Cue:LWin-Neut^*^, there is heightened estimated activity in the NAc, anterior insula and visual regions. Furthermore, task-relevant NAc activity is meaningfully higher in the Saturated model than the CueFeedback model for the Cue:LW-Neut^*^, Cue:LL-Neut^*^ and Cue:L-Neut contrasts.

Some MID task experts may view the potential for bias in Cue-based contrasts when Fixation is unmodeled in the CueFeedback model as less problematic due to its reflection of anticipatory neural signals. Given the previously mentioned differences in interpretations of the Cue- and Fixation-related signals, it is not clear whether allowing some unknown amount of Fixation signal to bias the Cue signal estimate is easily interpreted in the CueFeedback model. Large-scale studies like ABCD, which make activation estimates publicly available, are invaluable for advancing the field. However, the growing diversity of researchers using these data highlights the importance of providing contrast estimates with minimal ambiguity as those scientists may lack sufficient fMRI and task-specific training to make these distinctions about degrees of bias, instead assuming that the contrasts are truly valid measures of the labeled constructs.

### 4.1.4 Feedback comparisons with baseline highlight a modeling weakness

The activation maps for Feedback comparisons to baseline are broadly negative (Figure 5). We suspect this could be due to a poor fit of the canonical HRF for the Cue stimulus, which has previously been hypothesized to have a large undershoot (Srirangarajan et al., 2021). Unfortunately, investigating different HRF shapes would require fitting the HRF derivative and dispersion parameters or another basis set that can more flexibly capture the HRF shape. Further improvements to the Saturated model of ABCD’s MID task are constrained by the design of the task due to the previously mentioned collinearity where contrasts of interest had cVIFs exceeding 100. Addressing this issue would likely require a revised paradigm with greater temporal separation and jitter between events. One important consideration is that Fixation duration varies systematically with trial outcome (Hit vs. Miss), so any redesigned paradigm should account for this dependency when aiming to better characterize HRF shapes in the MID task. When designing such a paradigm, an efficiency/cVIF evaluation of the basis set models for the newly proposed study paradigms should be compared.

This bias is assumed to be the same for Hit and Miss trials within the same cue type, which protects the interpretation of Feedback-based Hit/Miss comparisons within Cue type, but Feedback comparisons within trial outcome between Cue types should be avoided, as discussed further below.

#### 4.1.5 Feedback-based outcome comparisons within condition

Given the similarity in efficiencies for FB:LHit-LMiss^*^ and FB:WHit-WMiss^*^ (Supplemental Figure S4), the activation differences found in our real data analyses (Supplemental Figure S14) likely reflect changes due to improvements in bias reduction with the Saturated model. Generally the activation is more negative for the Saturated model, which aligns with the positive bias introduced by Fixation and Probe activation found in our simulations for the CueFeedback model’s contrasts (Figure 4). Furthermore, there is higher activity in the insula for the within condition miss versus hit trials, illustrating the Saturated model is meaningfully better at capturing the valence activity during unsuccessful trials, most evidence for Loss trials.

However, questions remain about the interpretation of other Feedback-related condition comparison contrasts. Previously, contrasts comparing trial types within an outcome category (e.g., FB:WHit–NeutHit) were discouraged due to biases from unmodeled Fixation differences. While the Saturated model accounts for Fixation, the negative Feedback versus baseline activation discussed earlier, implies a potential mismodeling of the HRF and so Cue-based activation could still impact contrasts like FB:WHit-NeutHit. For Feedback-based contrasts within trial-type, like FB:LHit-LMiss^*^, under the assumption that the bias from the mismodeled Cue and Fixation is the same for all Loss trials, this bias would subtract out in this contrast. On the other hand, since Win and Neutral trials elicit very different magnitudes of Cue and Fixation activation, this bias does not subtract out in FB:WHit-NeutHit. Generally, FB contrasts versus baseline and FB contrasts comparing trial types should not be studied in either the CueFeedback or Saturated models, due to biasing potential.

### 4.2 Recommendations

It is common for a researcher to repeatedly use a task in multiple studies and some groups may have study designs that were previously found to be optimal in the efficiency analyses carried out when designing the historical studies. We recommend that moving forward, even if it is a task that a researcher has previously designed and worked with, unless the Saturated modeling approach was used, new efficiency analyses with the Saturated model should be carried out. We also recommend using our newly introduced cVIF calculation (Appendix C) to evaluate collinearity since it is possible that the range of designs studied in an efficiency analysis all have high collinearity and hence the most efficient design of that set will still have detrimental collinearity present.

If data have already been collected and the Saturated model for that task has high collinearity, it is often the case that events that are short in duration and adjacent will need to be combined as shown in Panel C of Figure 1. This is often the case for very short cues followed by very short response probes. This does limit the interpretation of the effect size estimates, but the only other alternative is to model the events without performing inferences on their activation estimates. This is a common problem encountered when first translating a behavioral study for use in fMRI as behavioral studies do not have the same timing limitations that are imposed in fMRI data due to the sluggish BOLD response that is impacted by hemodynamic delay.

## 5 Conclusion

The Saturated model, which includes all task events, is often approached with caution due to concerns about collinearity. We have introduced a more targeted tool, cVIF, for assessing the impact of collinearity on the estimates of interest: *contrast* estimates. Although the Saturated model has been dismissed as impractical for tasks like ABCD’s MID due to collinearity concerns, our cVIF tool and efficiency comparisons provide informed assurance that the Saturated model can be used comfortably with ABCD’s MID task. The Saturated model is preferred as it limits the potential biases quantified in our simulation study when using the CueFeedback model. However, improving the assumed HRF in the Saturated model is limited by the MID paradigm, which does not permit the use of basis sets needed to test alternative HRF shapes. Given this, some contrasts are still restricted from study when using the Saturated model with the MID task.

We recommend using both efficiency and cVIF when evaluating Saturated models during the design of task paradigms, as small timing changes can substantially reduce collinearity. When paradigm changes are not feasible, cVIF can help identify contrast estimates with high variability that may require further scrutiny. We also advise against the collinearity reduction strategies shown in Figure 1, as some fail to reduce collinearity while others compromise interpretation and/or introduce bias. If such strategies are used, their limitations should be clearly acknowledged.

## 6 Ethics

The ethical considerations for the ABCD data are presented in Clark et al. (2018).

## 7 Data and Code Availability

Code for the cVIF estimate can be found at https://github.com/jmumford/vif_contrasts. Code for the simulation analyses is publicly available (R. Poldrack & Mumford, 2025). Although the real data (event timings and response times) used to construct the simulated data cannot be distributed, we have supplied an alternative dataset that can be used to run the simulations within the code repository. The real data analysis code is also publicly available (M. Demidenko et al., 2025).

The ABCD BIDS data and fMRIPrep v23.1.4 derivatives can be accessed through the ABCD-BIDS Community Collection (ABCC) with an established Data Use Agreement (see https://abcdstudy.org/). The data used in these analyses will be available at a future release onto the NIH Brain Development Cohorts (NBDC) Biorepository Portal. The complete set of group-level maps are publicly available on Neurovault (84 images; https://identifiers.org/neurovault.collection:18697).

## 8 Author Contributions

JAM: Simulation study and manuscript writing. MID: Real data analysis and manuscript writing. JMB, CH, EJF, HPG, DJH, SMN, TDW: Provided feedback about ABCD MID implementation and manuscript revision. RAP: Simulation study, manuscript writing and supervision.

## 11 Acknowledgements

The authors would also like to thank the research participants and staff involved in data collection of the Adolescent Brain Cognitive Development (ABCD) Study data. The ABCD Study is a multisite, longitudinal study designed to recruit more than 10,000 children ages 9 and 10 and follow them over 10 years into early adulthood. Data used in the preparation of this article were obtained from the ABCD Study (https://abcdstudy.org), from the ABCD-BIDS Community Collection (ABCC). This is a multisite, longitudinal study designed to recruit more than 10,000 children age 9-10 and follow them over 10 years into early adulthood. The ABCD Study® is supported by the National Institutes of Health and additional federal partners under award numbers U01DA041048, U01DA050989, U01DA051016, U01DA041022, U01DA051018, U01DA051037, U01DA050987, U01DA041174, U01DA041106, U01DA041117, U01DA041028, U01DA041134, U01DA050988, U01DA051039, U01DA041156, U01DA041025, U01DA041120, U01DA051038, U01DA041148, U01DA041093, U01DA041089, U24DA041123, U24DA041147.

A full list of supporters is available at https://abcdstudy.org/federal-partners.html. A listing of participating sites and a complete listing of the study investigators can be found at https://abcdstudy.org/consortium_members/. ABCD consortium investigators designed and implemented the study and/or provided data but did not necessarily participate in the analysis or writing of this report. The authors thank Brian Knutson, Ryan Yan, and Joshua W. Buckholtz for helpful comments on earlier versions of this work.

## A Definitions of Efficiency and tVIF

For a regression model given by

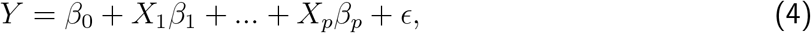

succinctly expressed in matrix form as *Y* = *Xβ* + *ϵ*, the variance of a contrast of parameter estimates is given by

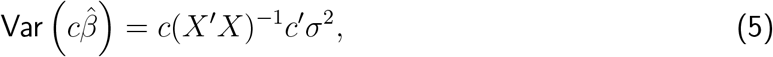

where *c* is a row vector of length *p* + 1 and *σ*^2^ is the residual variance. Efficiency ignores the unknown component of the variance, *σ*^2^, and focuses on the portion of the variance originating from the design matrix and is defined as

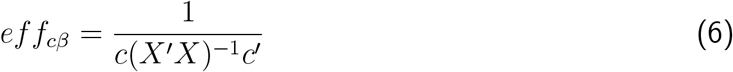

Friston et al. (1999) and Liu et al. (2001).

The traditional VIF (tVIF) is based on a ratio for the variance for a *β*_*j*_, Var (*β*_*j*_), in a model that only includes an intercept and *X*_*j*_ to the full model defined in Equation 4. If 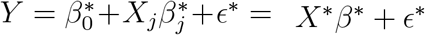 is this simplified model, the tVIF is given by

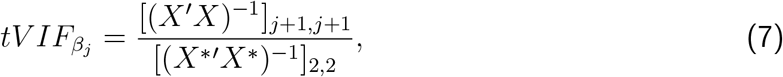

where [(*X*^*′*^*X*)^−1^]_*j*+1,*j*+1_ and [(*X*^*\*′*^*X*^*\**^)^−1^]_2,2_ are the diagonal elements corresponding to *X*_*j*_ (one is added due to the intercept). This view of tVIF clarifies that it is the ratio of the variance from the worst case scenario (all collinearity present) to the best case scenario (no collinearity). This is equivalent to the formulation presented in most regression texts, 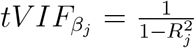, where 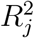 is the R-squared from the regression modeling *X*_*j*_ as a function of all other regressors, *X*_*j*_ = *β*_0_ + *X*_1_*β*_1_ + … + *X*_*j*−1_*β*_*j*−1_ + *X*_*j*+1_*β*_*j*+1_ + … *X*_*p*_*β*_*p*_ + *ϵ* (Neter et al., 1996)[Chapter 9].

## B How VIFS can be misleading

As shown in Appendix A, the traditional VIF (tVIF) is the ratio of the variance of a parameter estimate in the worse case scenario (collinearity present between regressors) to the best case scenario (no collinearity from other regressors). There are two shortcomings to this estimate of VIF. First, it cannot be used to estimate the VIF for contrasts of parameters. Second, the correlation between regressors may not reflect the true underlying correlation of the conditions as the example shown here illustrates.

The below example, also reflected in Panel A of Figure 1, illustrates how a correlation between two events, *A* and *B* can be masked from a traditional VIF calculation since the correlation between *A* and *B* is not captured by the correlation between the model regressors due to the use of orthogonal regressors. We then illustrate how our cVIF calculation captures the impact of correlation between conditions on the variance of the contrast.

We use a simple fMRI task with two conditions, A and B, and the goal is to test the condition difference. The models mirror the example in the first panel of Figure 1. Define Model 1 as,

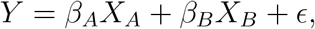

where *X*_*A*_ is a regressor that models condition A versus baseline and *X*_*B*_ models condition B versus baseline, and define Model 2 as,

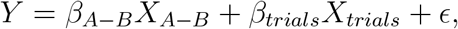

where *X*_*A*−*B*_ is a parametrically modulated regressor that models the difference between the A and B conditions coded using 1’s for A and -1’s for B (modulation values are then mean centered) and the *X*_*trials*_ regressor represents the average activation across all trials. Due to the centering of the modulation values, *X*_*trials*_ is orthogonal to *X*_*A*−*B*_ since the dot product of the modulation values of *X*_*A*−*B*_ (centered 1/-1 values) with the modulation values of *X*_*trials*_ (all 1s, since the regressor is not modulated) is 0. The test statistics for the A/B comparison will be identical from these two models, as such, the impact of condition correlations on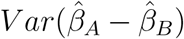from Model 1 will be the same as the impact on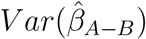 from Model 2, which is illustrated in the left panel of Figure A.1 for a range of condition correlations. Assume the regressors and outcome have been centered and scaled by the standard deviation/(N-1) where N is the number of observations. This obviates the need for an intercept in the model and ensures that the off-diagonal entry of the matrix *X*^*′*^*X* is the correlation between the two regressors in the model,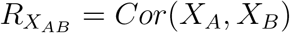. Note that these changes to the design matrix do not impact the inferences for the resulting parameter estimates (Neter et al., 1996, Chapter 7).

Focusing on Model 1, it follows that

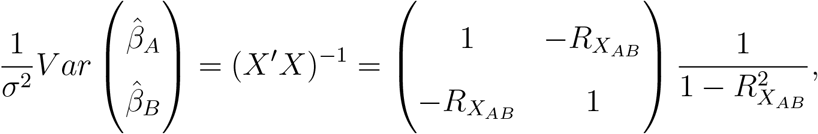

and so the worst case scenario variance, accounting for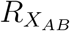, the correlation between the A and B conditions is,

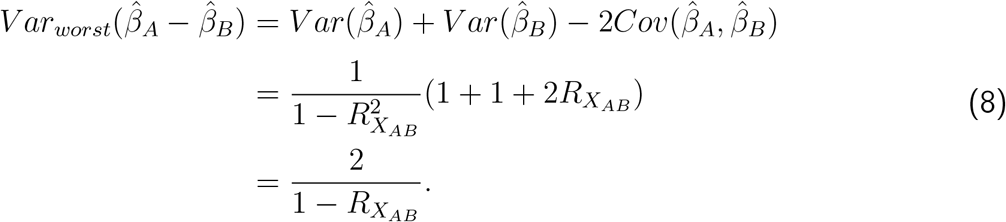

Our proposed cVIF calculation explicitly estimates the ratio of the contrast variance in the “worst” versus “best” case scenario. The “best-case” scenario for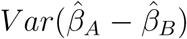 would then be if the conditions were not correlated which is when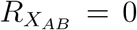. This yields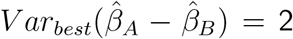, giving a contrast VIF of

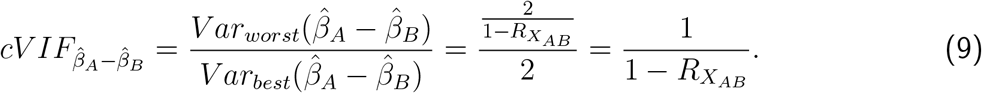

For Model 2, the traditional VIF for 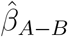, based on the the *R*^2^ from the regression modeling *X*_*A*−*B*_ as a function of *X*_*trials*_, takes on a value of 1 regardless of the correlation between the conditions since the regressors are uncorrelated and and do not reflect the correlations between conditions. Figure A.1 shows the relationship between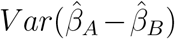 and the correlation between conditions, which is reflected in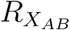, illustrating that only positive correlations between conditions has a negative impact on the condition difference. The right hand panel shows how the the cVIF estimate proposed here,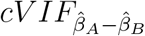mirrors the variance trend correctly indicating when the variance is inflated. The cVIFs from Model 1 corresponding to the individual condition effect estimates and are only helpful in identifying collinearity when conditions are positively correlated and can misidentify high collinearity when the conditions are negatively correlated. The traditional VIFs from Model 2 are uninformative. We recommend using the general cVIF calculation for contrasts defined in Appendix C along with the properly parameterized model.

## C cVIF: VIF calculation for contrasts of parameter estimates

A requirement for using this cVIF calculation for contrasts of parameter estimates is that each condition must be modeled within a separate regressor, as in Model 1 of the illustration in Appendix B. Parametrically modulated regressors should only be used if there are more than 2 levels for the modulator and in the case that there are only 2 levels, they can be equivalently expressed using separate regressors for each level. The design matrix entered into the calculation should omit the intercept, but mean center each regressor. This is an important step to ensure the correlations (not the dot products) between all pairs of regressors is set to 0 in the “best” case scenario. Additionally, it is useful to divide each regressor by its standard deviation as this may stabilize the estimate of the matrix inverse. Note that these changes to the design matrix do not affect the parameter estimates (Neter et al., 1996, Chapter 7). While centering and scaling are helpful for estimating cVIF, they are not necessary for the data analysis itself. In summary, *X* consists of centered and scaled regressors omitting the intercept term.

**Figure A.1:**
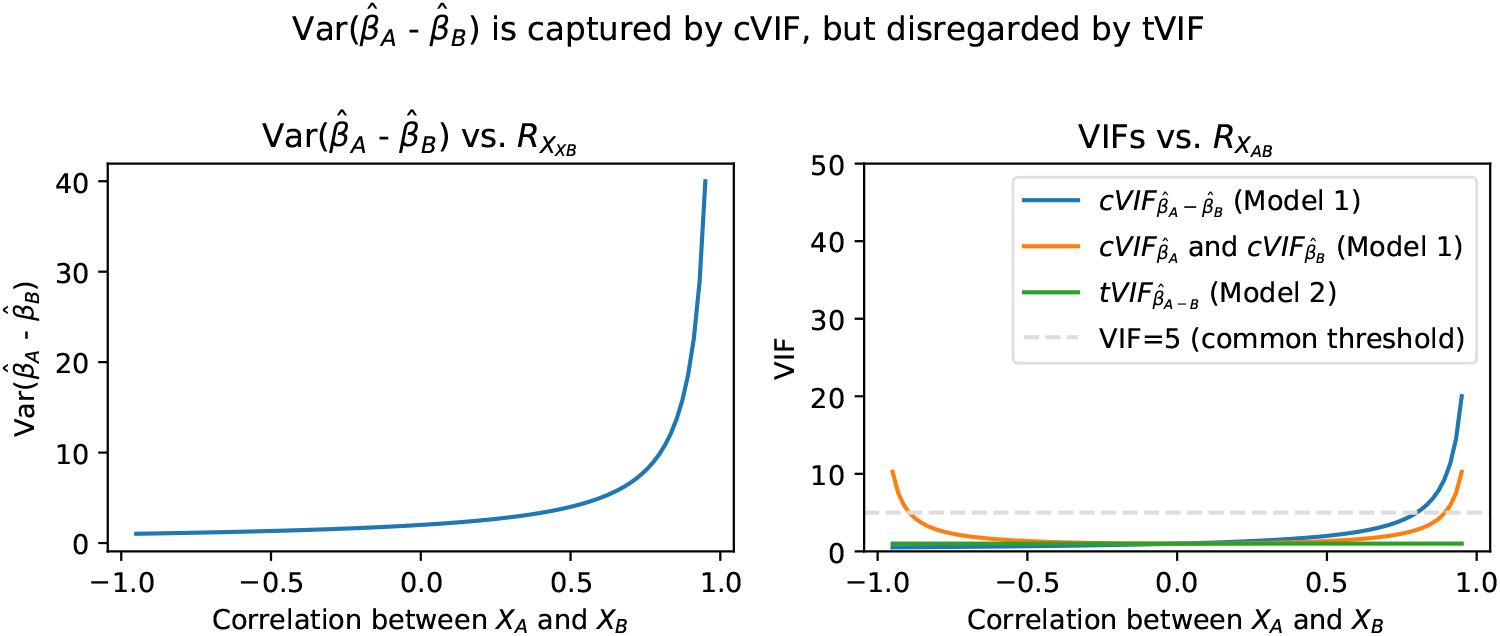
cVIF captures true impact of collinearity on the variance. The left panel shows how increasing positive correlations between conditions inflates the variance of the estimated contrast comparing conditions. The right panel compares our proposed contrast-based cVIF using Model 1 (blue, see Appendix C for the general algorithm) to using the traditional, condition-specific cVIFs from Model 1 (orange) and the traditional VIF from Model 2 (green), which is uninformative since it is constant due to its inability to reflect the correlation between the *A*/*B* conditions.

To motivate the calculation, for the linear regression specified by

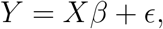

where *Y* is a vector (e.g., the BOLD time series), *X* is the design matrix (the centered regressors described above), *β* is a vector of parameters and *ϵ* is the error term, the variance of a contrast, row vector *c*, of parameter estimates is given by

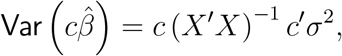

where *σ*^2^ is the residual variance. The cVIF estimate is a ratio of this variance under two conditions: the true condition correlations (the equation as given) versus no correlation between conditions (set all off-diagonal elements of *X*^*′*^*X* to zero). Note the residual variance term drops out in the ratio, so this calculation is not dependent upon the data, *Y*. Given this, general cVIF calculation is defined by

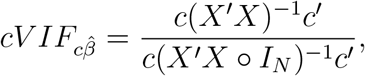

where the ○ operator represents the element-wise matrix multiplier and *I*_*N*_ is an *N* × *N* identity matrix, where *N* is the number of observations. The multiplication with *I*_*N*_ in the denominator sets the correlation between all conditions to 0.

A Python implementation of this algorithm can be found at https://github.com/jmumford/vif_contrasts.

