## Supplemental Materials for "Unintended bias in the pursuit of collinearity solutions in fMRI analysis"

#### S1 Accuracy of onset times in the ABCD data distribution

In May 2024, it was discovered and confirmed by the DAIRC that there was an error in the (a) Hagler et al. 2019 description of the number of calibration volumes in GE scanners and (b) the calculation of the behavioral timing files used in the task fMRI first level analyses and released with the minimally preprocessed data. First, in the initial description of the ABCD acquisition and task analysis protocol (Hagler et al., 2019), it was stated that the calibration sequence made up

16 volumes in the GE DV26, 5 in GE DV25 (one of the calibration volumes appears to comprise a collapsed 12 calibration volumes), 8 in Siemens and 8 in Philips scanners. It was inferred that the E-Prime tasks are triggered consistently across GE, Siemens and Philips scanners. However, it was discovered that the E-Prime tasks are triggered by the scanner at the start of the 5th and 16th calibration volume in the GE DV25 and GE DV26 scanners, respectively, instead of the start of the 6th and 17th volume). Meanwhile, in the Siemens/Philips scanners, the e-prime tasks are triggered correctly, starting following the end of the 8th calibration volume (i.e., start of 9th volume). As a result, the timing information in the GE data was displaced by 800 ms (1 volume). This was confirmed by analysis of the original and revised data using response-locked averaging, which showed that the offset was evident in the neural time course (Figure S1). Nonetheless, the DAIRC's reanalyses of the data following this correction resulted in only minor differences in estimated contrast beta weights for the CueFeedback model.

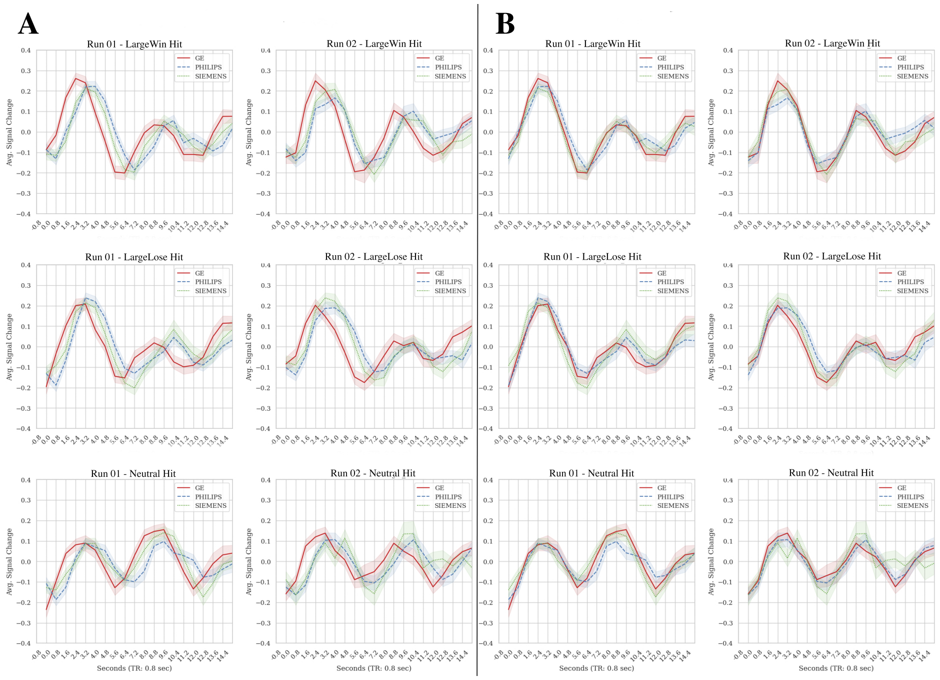


Figure S1: TR-by-TR plots locked to the **Probe onset** in the MID task for voxels averaged within an 8 mm radius sphere in the Left Motor Area (MNI: -38, -22, 56) for N = 50 GE, N = 50 Siemens and N = 50 Philips Baseline data for (A) Uncorrected and (B) Corrected data

### S2 Design Matrices in Simulations and Data Analyses

Both the simulated and real data analyses used the design matrices shown in Figure S2, but the real data analyses included additional nuisance regressors, such as motion regressors and discrete cosine filter with 128s cut-off.

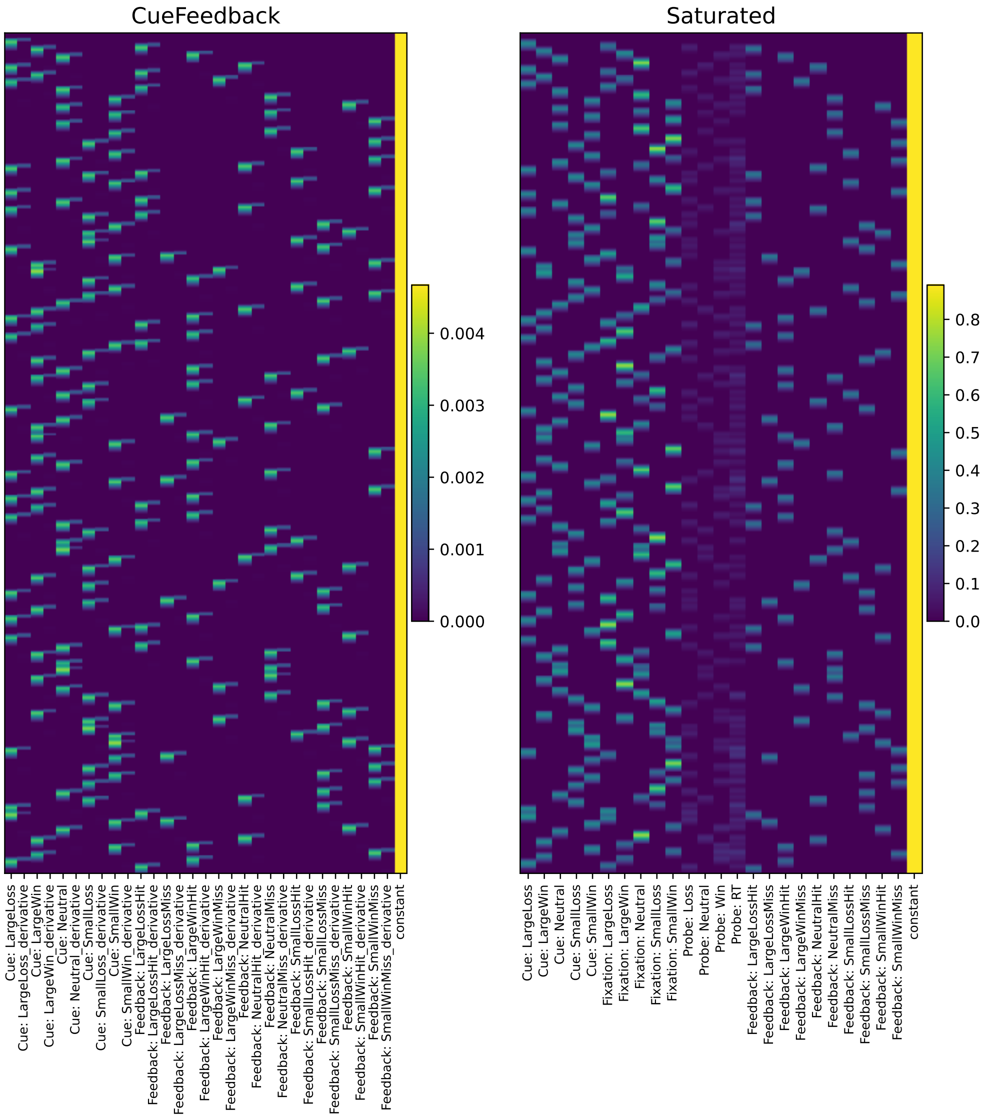


Figure S2: Design matrices used in simulated and real data analyses. The real data analysis design matrices also included cosine regressors to model low frequency drift (<0.008 Hz) and 12 motion parameters (translations, rotations and their derivatives).

### S3 cVIFs and efficiencies of design matrices

The cVIFs for all regressors and contrasts used in the simulation analyses are displayed in Figure S3 and the efficiencies for the regressors that are shared between the models and the contrasts are shown in Figure S4

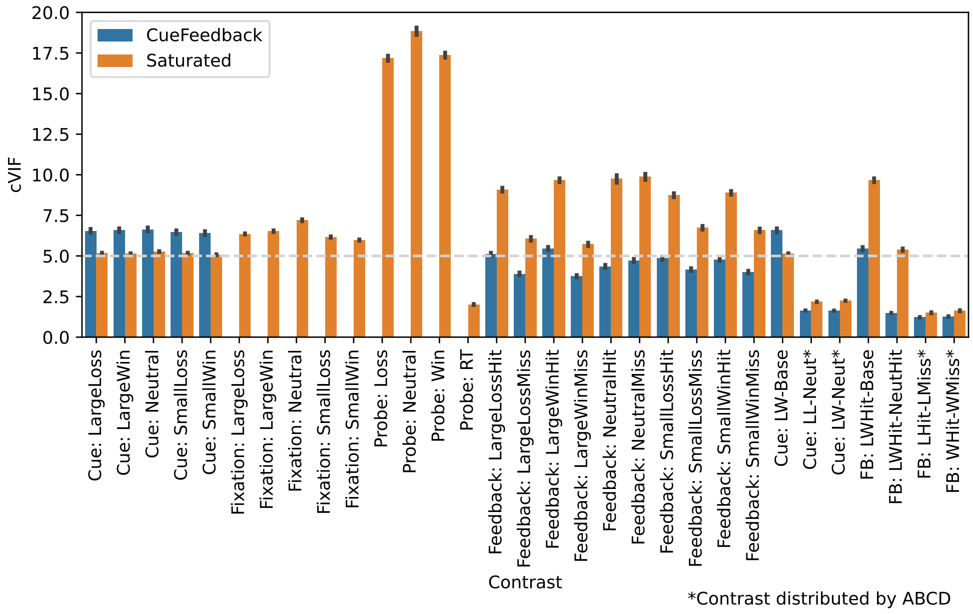


Figure S3: Variance inflation factors, using the cVIF algorithm, for all regressors and contrasts for both models.

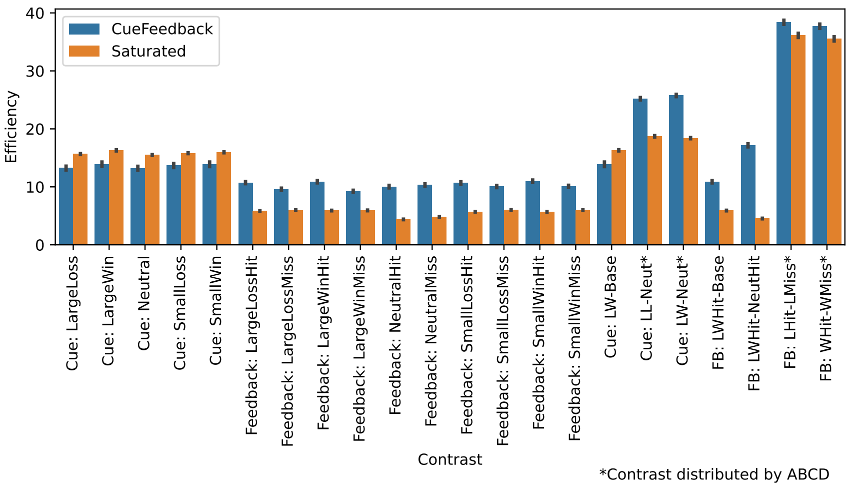


Figure S4: Efficiency estimates for all regressors that are shared by both models as well as the contrasts used in the simulation study.

### S4 Event durations and response times split by all three trial outcomes

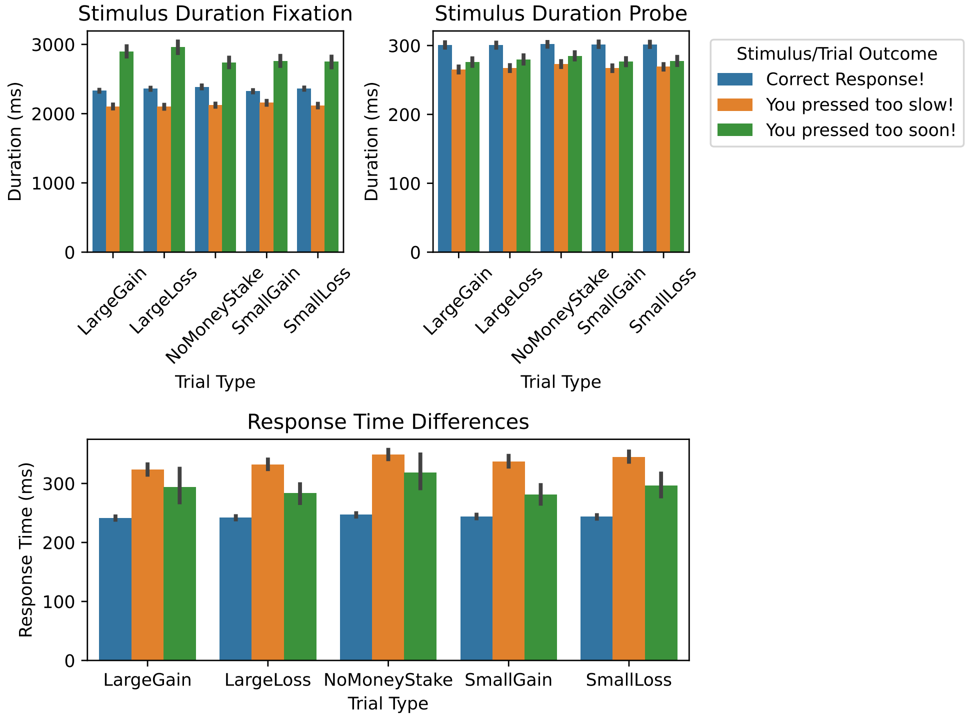


Figure S5: Event durations and response times split by Correct, Too Soon and Too Slow. Note that Miss trials consist of both Too Soon and Too Slow trials, which have very different durations and RTs. On average, 57%, 36% and 7% of trials are Correct, Too Slow and Too Soon, respectively.

### S5 Error rates and bias for all parameters and contrasts

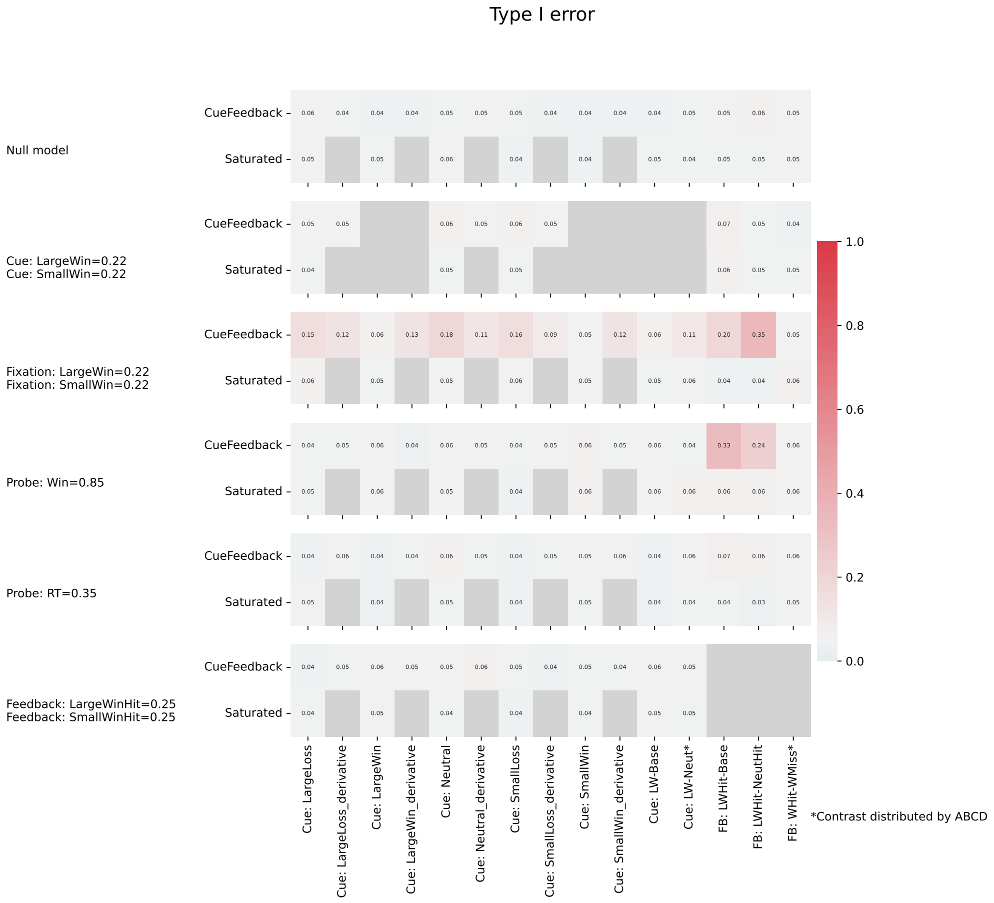


Figure S6: Type I error rates across simulation settings and models. Each row corresponds to a different model where the parameter or contrast error rate is shown for the ABCD (top) and Saturated (bottom) models, where values less than 0.05 indicate controlled Type I errors. Note the grayed out rectangles indicate that a parameter involved in that contrast was nonzero and so error rate is not estimated.

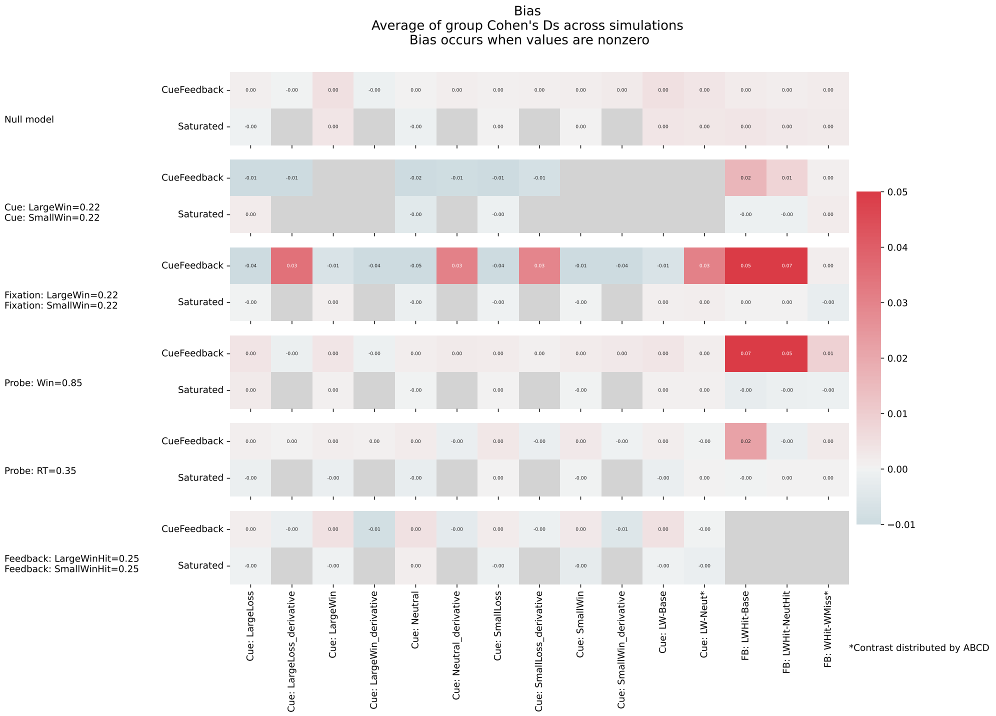


Figure S7: Bias in parameter and contrast estimates across simulation settings and models. Each row corresponds to a different model where the contrast estimate bias is shown for the ABCD (top) and Saturated (bottom) models, where 0 would indicate no bias. Note the grayed out rectangles indicate that a parameter involved in that contrast was nonzero and so bias is not estimated here.

### S6 Region of Interest: Nucleus Accumbens

Binarized masks for the Nucleus Accumbens (NAc) region of interest are created using the Harvard-Oxford subcortical atlases from *Nilearn*. First, the relevant subcortical regions— left and right NAc —are extracted based on their index values in the atlas. The ROI images are then resampled to align with the space of the reference beta map (MNI152 2009cAsym). The resampled ROI images are binarized to create binary masks Figure S8.

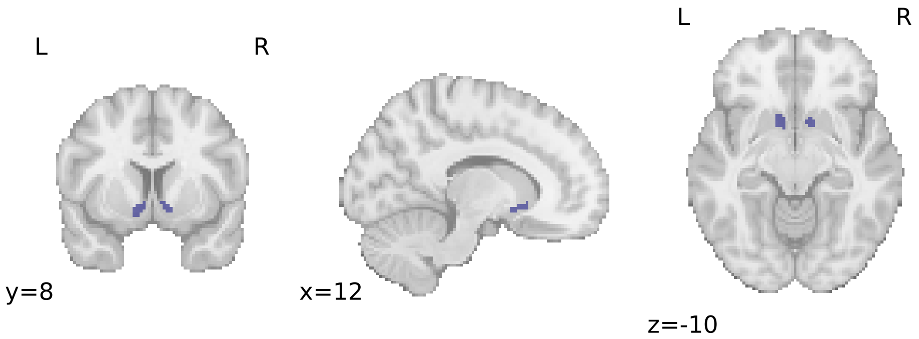


Figure S8: Harvard-Oxford subcortical Left and Right Nucleus Accumbens.

Differential NAc activation patterns between CueFeedback and Saturated models were evaluated by computing run-averaged statistical maps for each contrast condition. For each subject (N = 500), z-statistical maps were generated by dividing the beta contrast maps by the square root of the associated variance estimates. The resulting subject-level z-statistic maps were concatenated separately for the CueFeedback and Saturated models. The NAc masks were applied to the model specific concatenated z-statistic maps. For each contrast, the average z-statistic values within both right and left NAc ROIs were extracted. Paired difference plots were generated to visualize the model-dependent differences across each contrast condition within these task-relevant regions.

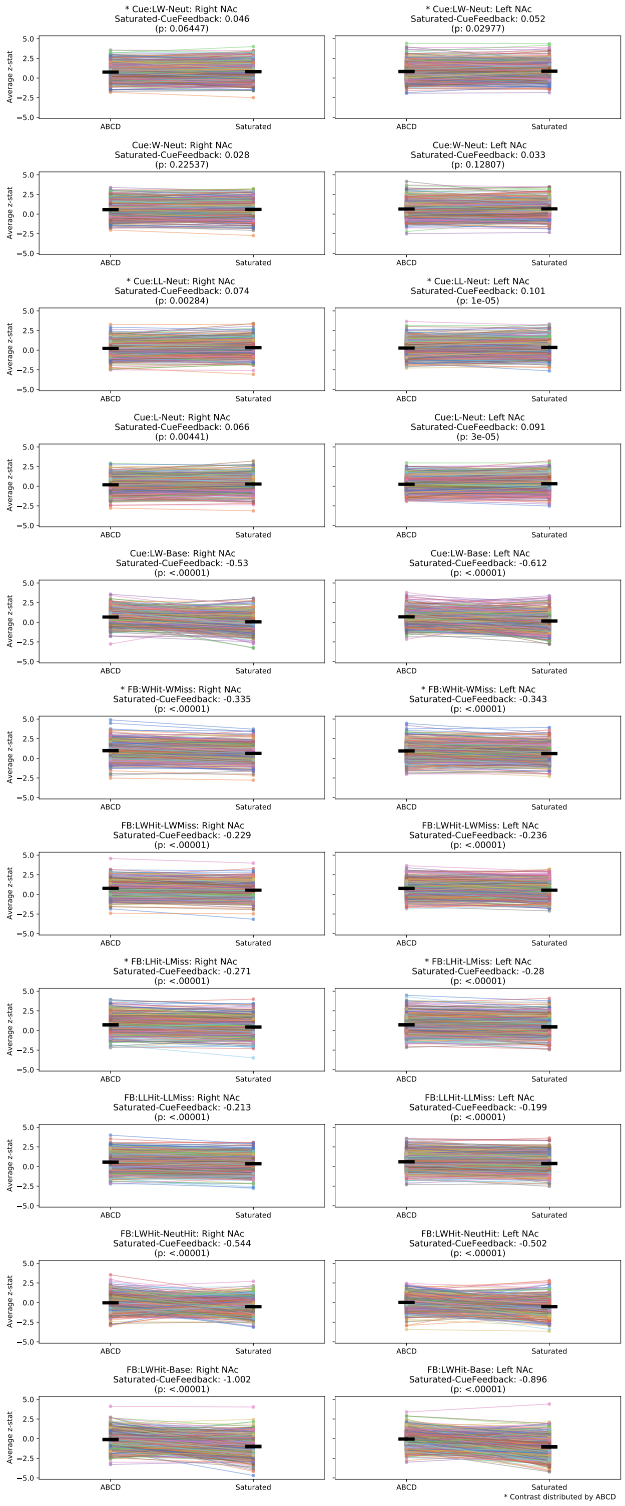


Figure S9: Paired difference plot for subjects' average z-statistical estimates withint he Left and Right Nucleus Accumbens for the CueFeedback and Saturated models. The difference between the Saturated − CueFeedback model and the associated *p*-value for that difference is reported for each contrast. * Contrasts included with ABCD data releases Chaarani et al., 2021.

### S7 Supplemental Behavioral Results

The average Probe hits (%) across Cue types are reported in Figure S10.

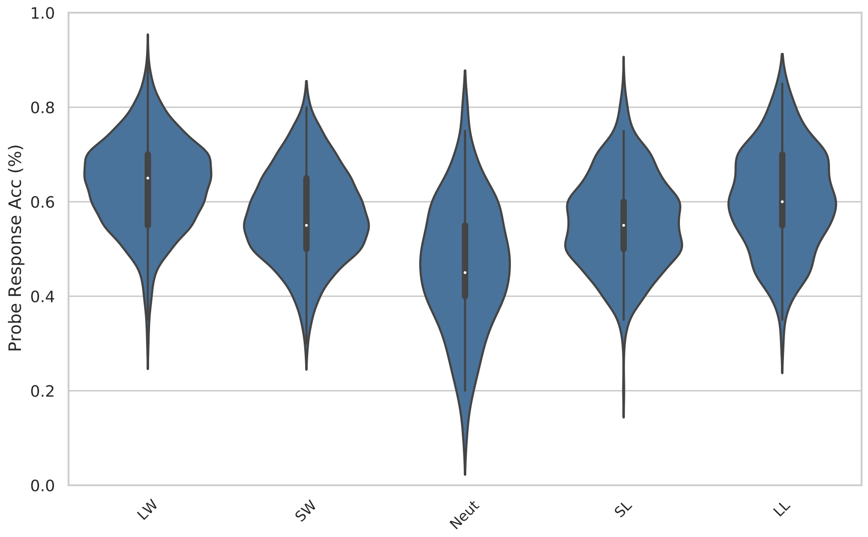


Figure S10: Probe response accuracy (%) across Cue type.

### S8 Supplemental Group-level Statistical Maps

In addition to the contrasts in the main text, we provide an expanded list of contrasts below in Table S1

| Contrast Name | Contrast Definition |
|---|---|
| **Cue: LW-Neut*** | 1 × Cue Large Win<br>- 1 × Cue Neutral |
| **Cue: W-Neut** | .5 × Cue Large Win<br>+ .5 × Cue Small Win<br>- 1 × Cue Neutral |
| **Cue: LL-Neut*** | 1 × Cue Large Loss<br>- 1 × Cue Neutral |
| **Cue: L-Neut** | .5 × Cue Large Loss<br>+ .5 × Cue Small Loss<br>- 1 × Cue Neutral |
| **Cue: LW-Base** | 1 × Cue Large Win |
| **Fix: LW-Neut** | 1 × Fixation Large Win<br>- 1 × Fixation Neutral |
| **Fix: W-Neut** | .5 × Fixation Large Win<br>+ .5 × Fixation Small Win<br>- 1 × Fixation Neutral |
| **Fix: LL-Neut** | 1 × Fixation Large Loss<br>- 1 × Fixation Neutral |
| **Fix: L-Neut** | .5 × Fixation Large Loss<br>+ .5 × Fixation Small Loss<br>- 1 × Fixation Neutral |
| **Fix: LW-Base** | 1 × Fixation Large Win |
| **Probe: All-Base** | .33 × Probe Win<br>+ .33 × Probe Loss<br>+ .33 × Probe Neutral |
| **Probe: Win-Loss** | 1 × Probe Win<br>- 1 × Probe Neutral |
| **Probe: Loss-Neut** | 1 × Probe Loss<br>- 1 × Probe Neutral |
| **Probe: Win-Neut** | 1 × Probe Win<br>- 1 × Probe Neutral |
| **RT** | 1 × RT |
| **FB: WHit-WMiss*** | .5 × Feedback Large Win Hit<br>+ .5 × Feedback Small Win Hit<br>- .5 × Feedback Large Win Miss<br>- .5 × Feedback Small Win Miss |
| **FB: LWHit-LWMiss** | 1 × Feedback Large Win Hit<br>- 1 × Feedback Large Win Miss |
| **FB: LHit-LMiss*** | .5 × Feedback Large Loss Hit<br>+ .5 × Feedback Small Loss Hit<br>- .5 × Feedback Large Loss Miss<br>- .5 × Feedback Small Loss Miss |
| **FB: LLHit-LLMiss** | 1 × Feedback Large Loss Hit<br>- 1 × Feedback Large Loss Miss |
| **FB: LWHit-NeutHit** | 1 × Feedback Large Win Hit<br>- 1 × Feedback Neutral Hit |
| **FB: LWHit-Base** | 1 × Feedback Large Win Hit |

* Contrast distributed by ABCD

Table S1: Definitions of contrasts focused on for our main real data analysis results.

#### S8.1 Cluster Corrected Statistical Maps

As mentioned in the main text, contrasts 1-4 and 11-14 are similar to those that are estimated and distributed in the ABCD consortium task fMRI derivatives Chaarani et al., 2021.

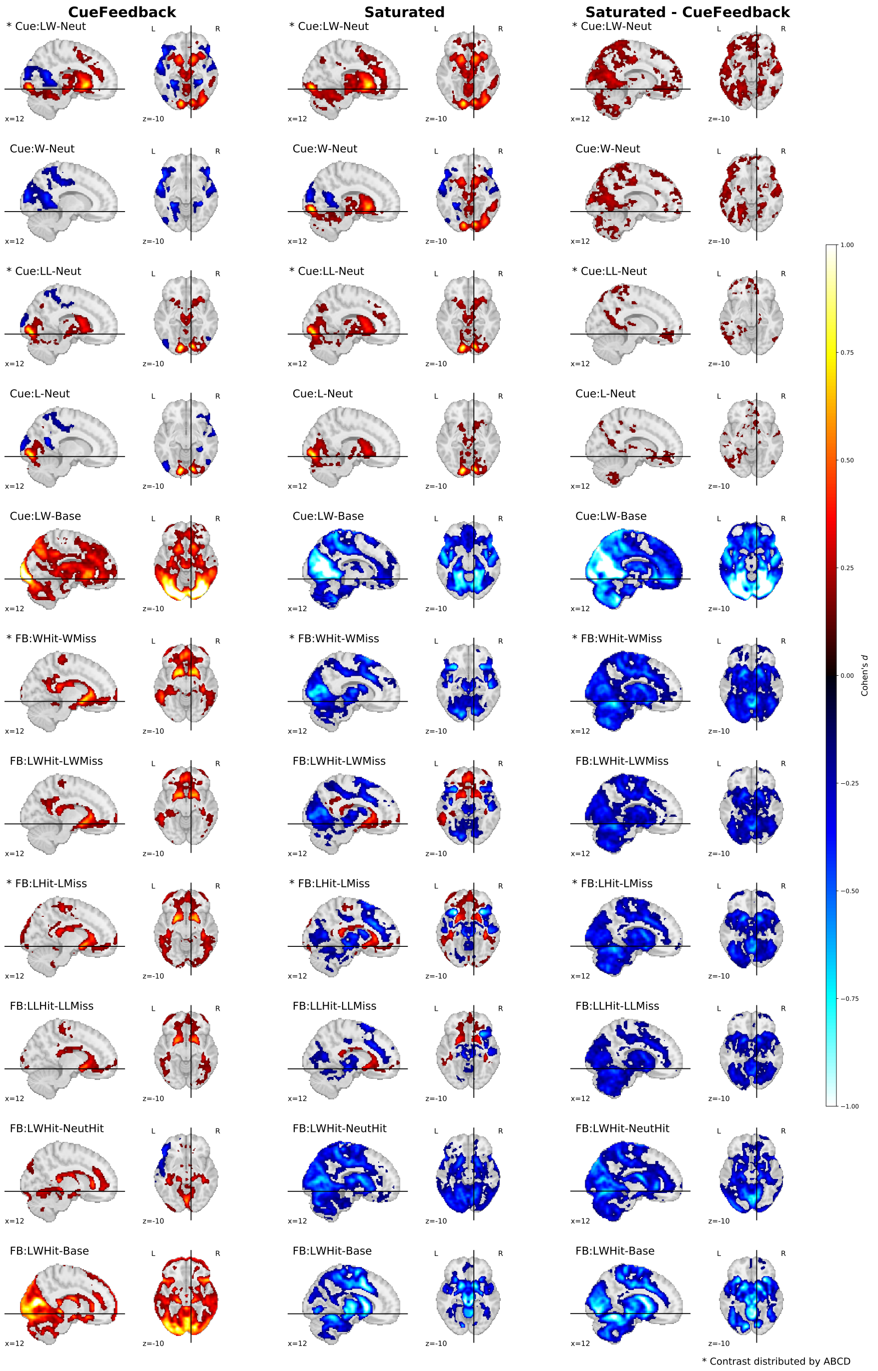


Figure S11: Cohen's *d* group-level activation maps (voxels selected using a cluster based permutation accounting for site, 8192 permutations) across Cue (5) and Feedback (6) contrasts for the CueFeedback (shown on the left), Saturated (shown in the middle) and their paired difference (shown on the right). We note contrasts included with ABCD data releases Chaarani et al., 2021.

Figure S12 illustrates the group-level effect of the Probe events and the Probe RT.

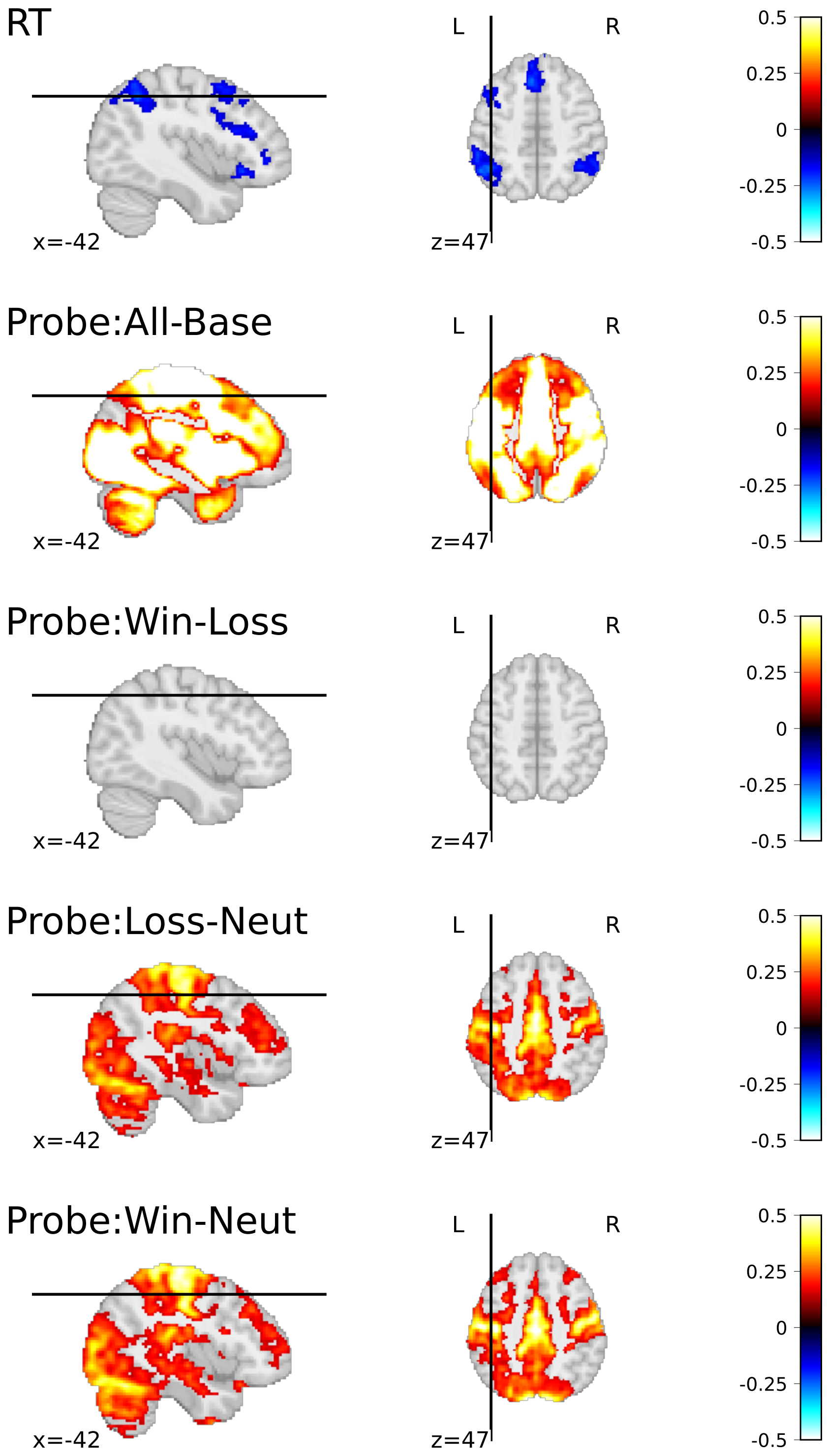


Figure S12: Group average effect (Cohen's $d$) of RT, Probe versus baseline and pairwise contrasts between Win, Lose and Neutral probe events (voxels selected using a cluster based permutation accounting for site)

Figure S13 presents the group-level activity for five contrasts across the Cue and Fixation components of the MID task trials.

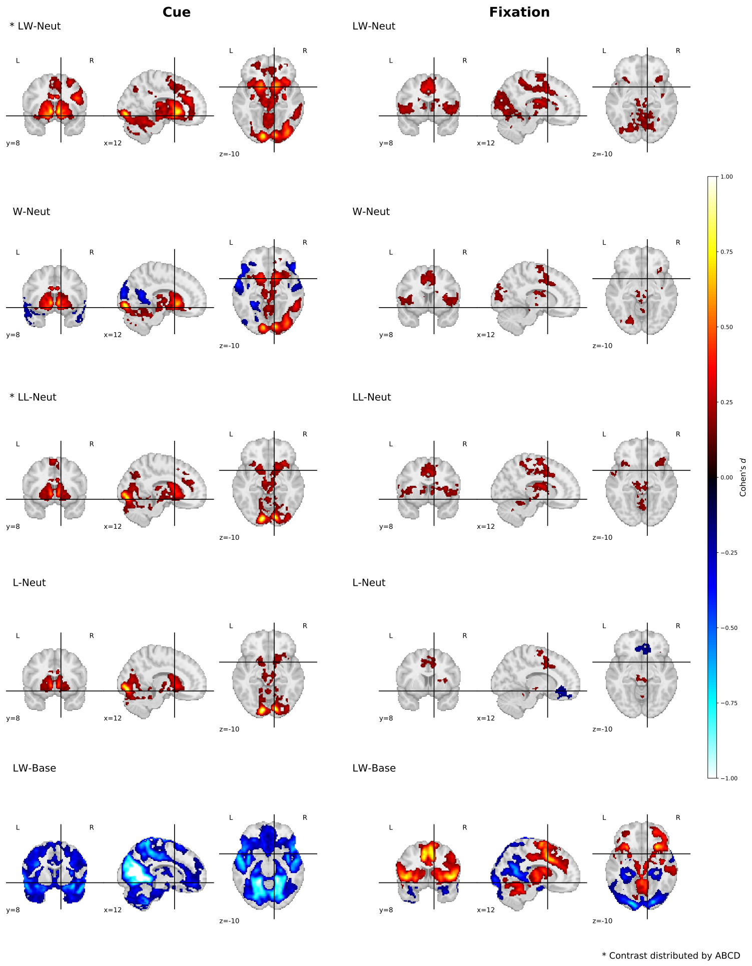


Figure S13: Cohen's *d* group-level activation maps (voxels selected voxels selected using a cluster based permutation accounting for site) across a Cue (5; shown on the left) and Fixation (5; shown on the right) contrasts for the Saturated model. We note contrasts included with ABCD data releases Chaarani et al., 2021.

#### S8.2 Cluster Corrected Statistical Maps

Similar to Figure 5, Figure S14 are cluster corrected and thresholded Cohen's *d* statistical maps.

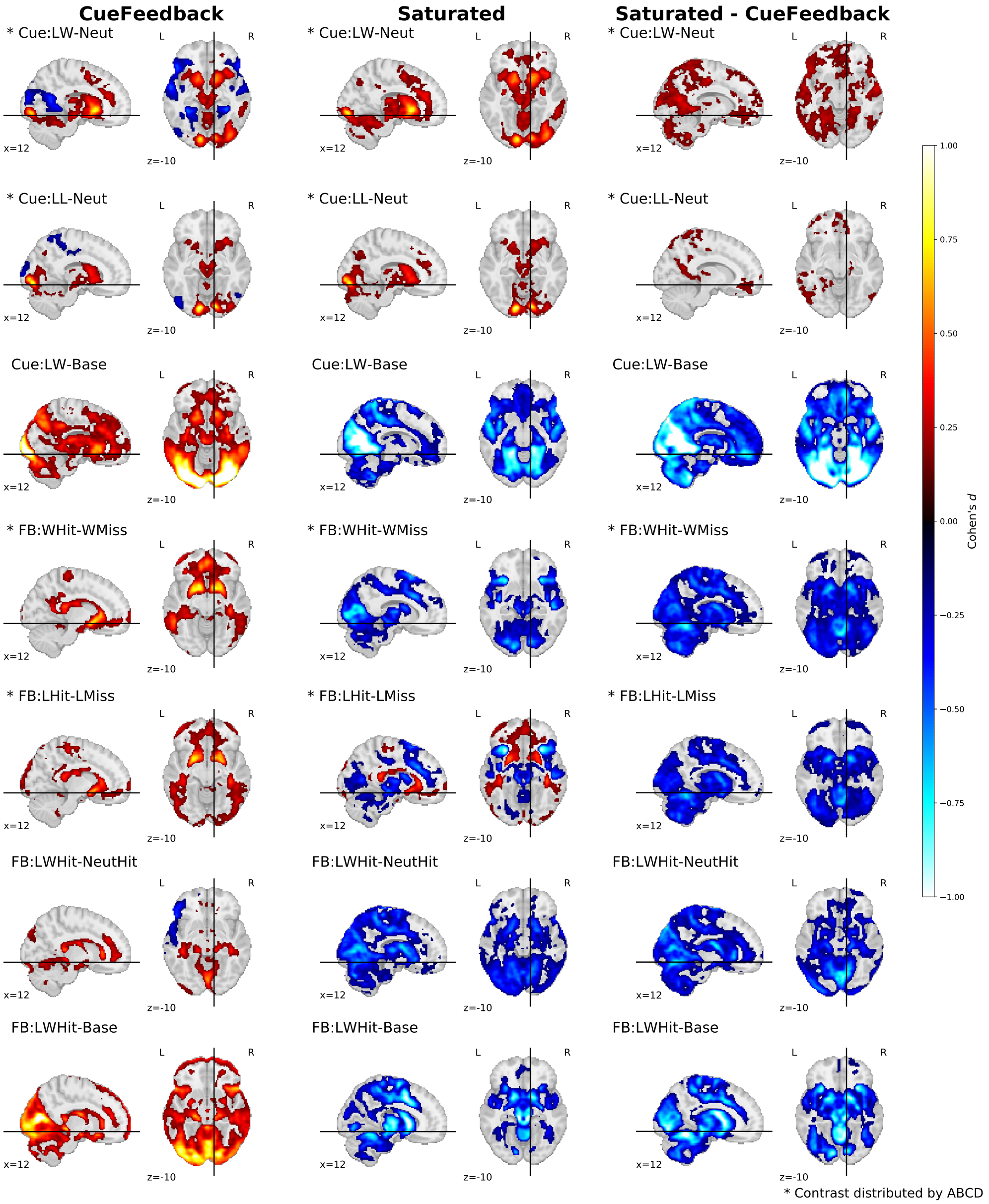


Figure S14: Cohen's *d* group-level activation maps (voxels selected using a cluster based permutation accounting for site, with 8192 permutations) across a subset Cue (3/5) and Feedback (4/6) contrasts for the CueFeedback (shown on the left) and Saturated model (shown in the middle) and the Paired statistical difference (Saturated−CueFeedback; shown on the Right). * Contrasts included with ABCD data releases Chaarani et al., 2021.

Similar to figure S13, the below figure S15 are non-cluster corrected and unthresholded Cohen's *d* statistical maps.

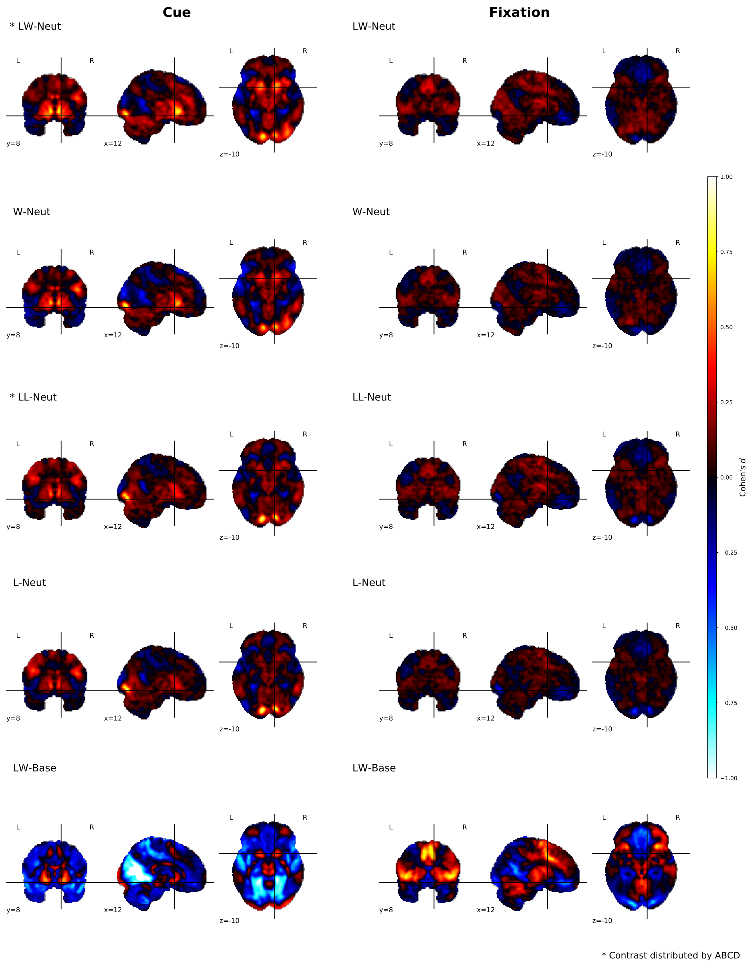


Figure S15: Cohen's *d* [uncorrected] group-level activation maps across a Cue (5; shown on the left) and Fixation (5; shown on the right) contrasts for the Saturated model. * Contrasts included with ABCD data releases Chaarani et al., 2021.
